# SAYSD1 senses UFMylated ribosome to safeguard co-translational protein translocation at the endoplasmic reticulum

**DOI:** 10.1101/2022.12.05.519155

**Authors:** Lihui Wang, Yue Xu, Sijung Yun, Quan Yuan, Prasanna Satpute-Krishnan, Yihong Ye

**Affiliations:** Laboratory of Molecular Biology, National Institute of Diabetes and Digestive and Kidney Diseases, National Institutes of Health, Bethesda, MD 20892; Department of Biochemistry, Uniformed Services University of the Health Sciences, Bethesda, MD 20814; Dendrite Morphogenesis and Plasticity Unit, National Institute of Neurological Disorders and Stroke, National Institutes of Health, Bethesda, MD, USA

**Keywords:** Co-translational protein translocation, Translocation-associated protein quality control/TAQC, SAYSD1, UFM1/UFMylation, Sec61, ribosome stalling/translation arrest, translocon clogging, collagen biogenesis

## Abstract

Translocon clogging at the endoplasmic reticulum (ER) as a result of translation stalling triggers ribosome UFMylation, activating a <u>T</u>ranslocation-<u>A</u>ssociated <u>Q</u>uality <u>C</u>ontrol (TAQC) mechanism that degrades clogged substrates. How cells sense ribosome UFMylation to initiate TAQC is unclear. Here we use a genome-wide CRISPR/Cas9 screen to identify an uncharacterized membrane protein named SAYSD1 that facilitates TAQC. SAYSD1 associates with the Sec61 translocon, and also recognizes both ribosome and UFM1 directly, engaging a stalled nascent chain to ensure its transport via the TRAPP complex to lysosomes for degradation. Like UFM1 deficiency, SAYSD1 depletion causes the accumulation of translocation-stalled proteins at the ER and triggers ER stress. Importantly, disrupting UFM1- and SAYSD1-dependent TAQC in *Drosophila* leads to intracellular accumulation of translocation-stalled collagens, defective collagen deposition, abnormal basement membranes, and reduced stress tolerance. Together, our data support a model that SAYSD1 acts as a UFM1 sensor that collaborates with ribosome UFMylation at the site of clogged translocon, safeguarding ER homeostasis during animal development.

## Introduction

In eukaryotes, proteins of the secretory pathway are synthesized by endoplasmic reticulum (ER)- bound ribosomes and inserted into the ER lumen or integrated into the membrane co-translationally through the Sec61 translocon (Rapoport et al., 2017). This process is highly sensitive to perturbations by translation stalling or defects in protein modification, folding, and assembly; all of which can generate faulty polypeptides that clog the translocon (Phillips and Miller, 2020; Wang and Ye, 2020). Failure to clear clogged translocons prevents nascent polypeptides from entering the ER, posing great harms to cells (Wang and Ye, 2020). While polypeptides stalled during post-translational translocation are cleared by a recently identified metalloprotease Ste24 (Ast et al., 2016), not much is known about how cells cope with polypeptides stalled during co-translational protein translocation, the major protein translocation pathway at the ER.

UFM1 is a small protein structurally homologous to ubiquitin. Like ubiquitin, it can be conjugated to target proteins through UFMylation, a process mediated by an enzyme cascade involving UBA5 (E1), UFC1 (E2), and an ER-localized trimeric ligase complex (E3) composed of UFL1, DDRGK1 (also named UFBP1) and CDK5RAP3 (Gerakis et al., 2019). The reaction is reversible as UFM1 can be cleaved off from substrates by two deUFMylating enzymes UFSP1 and UFSP2 (Kang et al., 2007). Genetic inactivation of UFM1 or UFMylating enzymes are known to cause ER stress (Cai et al., 2015; Lemaire et al., 2011; Zhang et al., 2012) and erythropoiesis defects (Cai et al., 2015; Tatsumi et al., 2011), whereas mutations in *UFM1* or genes encoding UFMylating enzymes have been linked to a variety of neurological disorders such as hypomyelinating leukodystrophy and epileptic encephalopathy (Colin et al., 2016; Duan et al., 2016; Muona et al., 2016; Nahorski et al., 2018; Gerakis et al., 2019).

We and others recently established the ribosomal subunit RPL26 as the principal target of UFMylation (Walczak et al., 2019; Wang et al., 2020). Importantly, we showed that RPL26 UFMylation occurs on the ER membrane and is specifically activated by co-translational translocation stalling (Wang et al., 2020). Ribosome UFMylation enables the export of stalled substrates from the ER for lysosomal degradation, suggesting a translocation-associated quality control (TAQC) mechanism that is distinct from the previously established ER-associated protein degradation (ERAD) (Sun and Brodsky, 2019) or ribosome-associated quality control (RQC) (Joazeiro, 2017). Besides ribosomes, UFM1 may modify other proteins at lower levels to regulate other cellular processes such as ERphagy and DNA damage response (Liang et al., 2020; Qin et al., 2019; Stephani et al., 2021).

To dissect how ribosome UFMylation promotes ER protein homeostasis, we conducted a genome wide CRISPR/Cas9 knockout (KO) screen using a genetically engineered translocon-clogging reporter. The identified genes collectively reconstruct a ‘translocon-declogging’ system centered around the UFMylation system, ribosome, the Sec61 translocon, and a novel translocon-associated membrane protein SAYSD1. We show that SAYSD1 binds UFMylated ribosome in a bipartite manner to facilitate the elimination of translation-stalled ER proteins. Importantly, our study identified endogenous collagens as a class of translation stalling prone protein and revealed a critical role for SAYSD1 and UFM1 in safeguarding collagen biogenesis during animal development.

## RESULTS

### Translocation-stalled nascent chains are eliminated by ER-to-lysosome transport

To study TAQC, we used a GFP-based model reporter, ER_GFP__K20 (Figure 1A) (Wang et al., 2020), which contains a signal sequence (SS) for co-translational ER targeting, a *N*-glycosylation site (-CHO) followed by GFP. Importantly, ER_GFP__K20 contains 20 consecutive lysine residues encoded by a polyadenine (poly-A) segment known to stall ribosomes during elongation (Ito-Harashima et al., 2007; Juszkiewicz and Hegde, 2017). It also contains mCherry (mCh) downstream of the stalling sequence whose expression would indicate translation read-through. We showed previously that translation-stalled ER_GFP__K20 is degraded by lysosomes following UFM1-dependent export from the ER (Wang et al., 2020).

**Figure 1.**
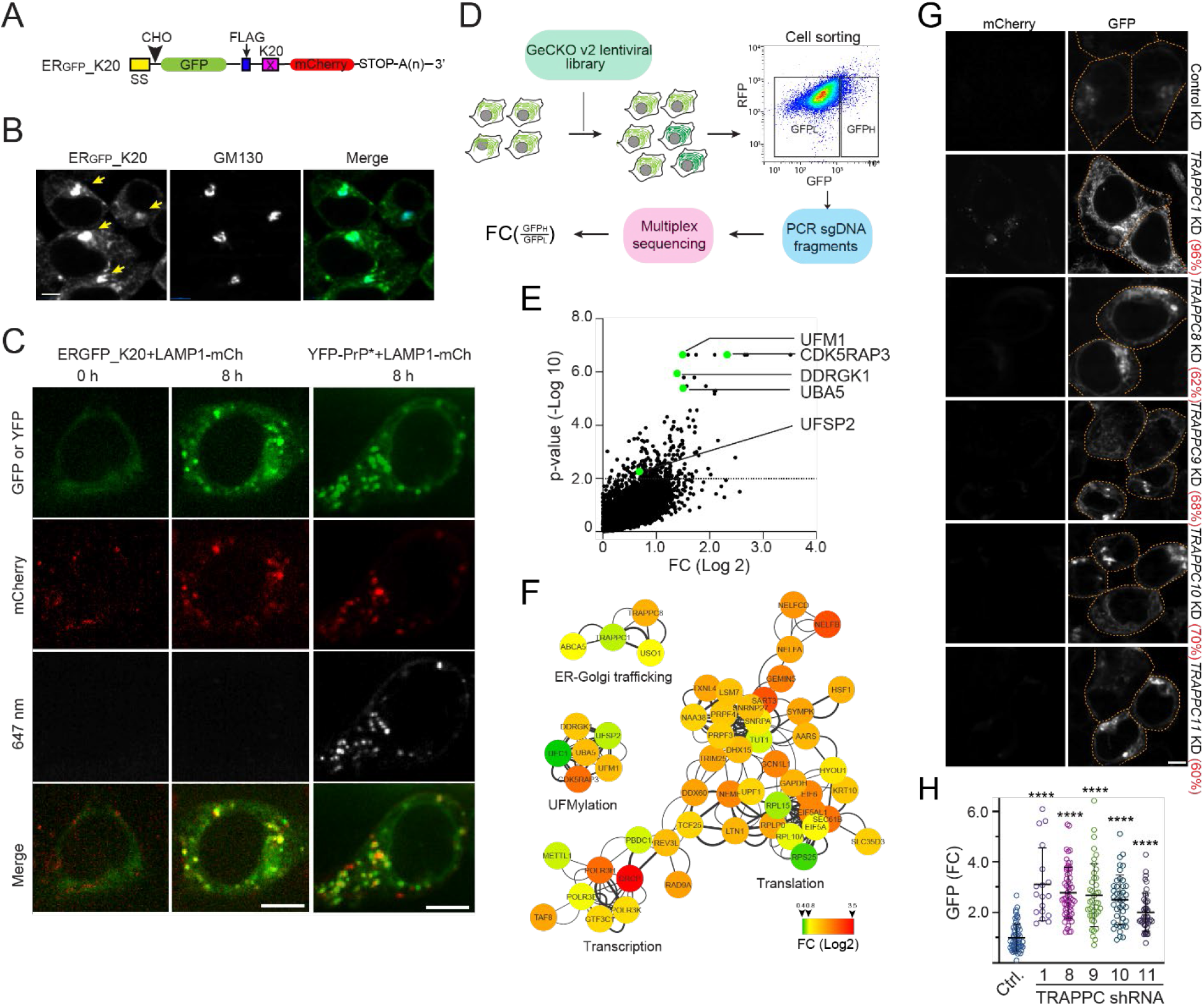
ER_GFP__K20 is transported to lysosome via the Golgi by the TRAPP complex. **(A)** A schematic diagram of the model TAQC substrate. SS, signal sequence; CHO, *N*-glycosylation site; X=K20. **(B)** ER_GFP__K20 is exported to lysosomes via the Golgi. ERGFP_K20-expressing 293T cells were treated with Bafilomycin A1 (Baf A1) (200 nM), fixed, and stained by GM130 antibodies (blue). Arrows indicate Golgi-localized ER_GFP__K20. Scale bar, 5 μm. **(C)** Representative images from time-lapse videos show that ER_GFP__K20 and YFP-PrP* are transported to lysosomes via distinct routes. ER_GFP__K20 stable cells transfected with LAMP1-mCherry (mCh) were imaged at the indicated time points after treatment with Baf A1 and Alexa_647_-labeled GFP antibodies (the left panels). The right panels show cells transfected with YFP-PrP* and LAMP1-mCh as a control. **(D)** A schematic diagram of the CRISPR/Cas9 screen. LFC, Log fold change. **(E)** A scatter plot shows the distribution of genes positively enriched in ER_GFP__K20 high cells. The dotted line indicates *p*= 0.01. Note that sgRNAs targeting the UFM1 pathway are enriched. **(F)** A Cytoscape gene interaction map for sgRNAs positively enriched in ER_GFP__K20 high cells. The color key indicates fold change (FC) in Log 2. **(G, H)** Knockdown of TRAPPC genes causes ER_GFP__K20 accumulation in cells. **(G)** Representative confocal images showing ER_GFP__K20 stable cells transfected with the indicated shRNA constructs. The numbers indicate average knockdown efficiency determined by qRT-PCR (n=3). **(H)** The graphs show the quantification of GFP fluorescence in individual cells. Error bars indicate means ± SD; **** *p*<0.0001 by one-way ANOVA with Dunnett’s multiple comparisons test. n=3 independent experiments.

To understand how ER_GFP__K20 is transported to lysosomes, we performed live-cell imaging using ER_GFP__K20 stable cells treated with the lysosome inhibitor Bafilomycin A1 (Baf A1). The result showed that after treatment, ER_GFP__K20 was first accumulated in a perinuclear region before reaching the lysosomes (Supplemental video 1). Immunostaining showed an extensive colocalization of perinuclear ER_GFP__K20 with the Golgi protein GM130 (Figure 1B). This observation, together with the finding that the degradation of ER_GFP__K20 is sensitive to the Golgi-disrupting agent Brefeldin A (Wang et al., 2020), suggests that trafficking through the Golgi is essential for lysosomal degradation of ER_GFP__K20.

The perinuclear trafficking of ER_GFP__K20 is reminiscent of rapid ER stress-induced export (RESET) of misfolded glycosylphosphatidylinositol (GPI)-anchored proteins (Satpute-Krishnan et al., 2014). Because RESET substrates are first exported to the plasma membrane before entering the endolysosomal system (Satpute-Krishnan et al., 2014), we tested whether ER_GFP__K20 followed the same trafficking itinerary. We incubated ER_GFP__K20 cells expressing LAMP1-mCh with Alexa_647_-labeled GFP antibody in the presence of Baf A1. If ER_GFP_-K20 was transiently exposed to the cell exterior, it should capture the GFP antibody, resulting in the co-internalization of ER_GFP__K20 with the antibody. As a positive control, we used cells expressing the RESET substrate YFP-tagged PrP*, which was also recognized by the GFP antibody (Satpute-Krishnan et al., 2014). As expected, YFP-PrP* cells accumulated GFP antibodies in LAMP1 positive lysosomes after incubation. By contrast, we failed to detect GFP antibodies in lysosomes of ERGFP-K20 cells under the same condition (Figures 1C, S1A-C), suggesting that unlike RESET, ER_GFP__K20 is transported via the ER-Golgi-lysosome path without going through the plasma membrane.

In UFM1 deficient cells, stabilized ER_GFP_-K20 is accumulated in the ER (Wang et al., 2020). To see if this phenotype results from a defect in releasing stalled substrates from the ribosome, we combined a pulse-chase assay with ribosome fractionation. S^35^-methionine labeled, newly synthesized ER_GFP_-K20 was immunoprecipitated from either the ribosome-bound or ribosome free fraction under a condition that preserved the peptidyl-tRNA linkage. As expected, autoradiography detected both tRNA-linked and tRNA-free ER_GFP_-K20 in ribosome associated fraction, but only tRNA-free ER_GFP_-K20 in the ribosome free fraction at the onset of the chase in both control and UFM1 knockdown cells (Figure S1D). In both cells, ribosome associated ER_GFP_-K20 species gradually disappeared during the chase due to substrate release, but only in UFM1 deficient cells, did we observed a consistent accumulation of ER_GFP_-K20 in the ribosome free fractions over time, suggesting that it is stabilized in UFM1-depleted cells after release from ribosomes. Since protease protection showed that ER_GFP_-K20 is entirely shielded by the ER membrane (Wang et al., 2020), we conclude that translation stalled ER_GFP_-K20 is released into the ER lumen in a UFMylation independent manner and then transported to the Golgi (see discussion).

### A genome wide CRISPR screen identifies TAQC regulators

To identify regulators of TAQC, we performed a genome-wide CRISPR-Cas9 knockout (KO) screen using 293T cells stably expressing ER_GFP__K20. We transduced cells with a lentivirus-based CRISPR/Cas9 library, targeting 19,050 human genes each with 6 sgRNAs (Sanjana et al., 2014). Cells were separated into GFP-high (10%) and GFP-low (85%) populations by fluorescence-activated cell sorting (Figure 1D). Next-generation sequencing of PCR-amplified sgRNAs identified those enriched in the GFP-high populations from two independent repeats, which correspond to 234 significant gene hits (*p* < 0.01) (Figure 1E, Table S1). STRING-based protein network analysis highlighted 88 high-confident hits with combined false discovery rate (FDR) below 0.05 (see methods). Most genes in this group are functionally interlinked, forming one large and two smaller clusters (Figure 1F). One small cluster includes most UFMylation genes such as *UFM1, UBA5, DDRGK1, CDK5RAP3* and *UFSP2* (Figures 1E, F), which validated the genetic screen. The large cluster includes many genes involve in transcription and mRNA splicing.

They might regulate ER_GFP__K20 at the mRNA level and thus were excluded from further studies. This cluster also includes genes involved in translation and ER protein translocation (e.g., ribosome subunits RPL15, RPLP0, RPL10A, RPS25, EIF6, EIF5A, and Sec61β). These proteins might have a dual role: regulating protein synthesis while ensuring proper ER translocation via TAQC. Interestingly, the screen also identified two ribosome-associated quality control (RQC) factors: Ltn1 and NEMF1. In RQC, NEMF1 recruits Ltn1 to 60S ribosome following ribosome splitting, resulting in ubiquitination of stalled substrates (Bengtson and Joazeiro, 2010; Shao et al., 2015). How precisely these factors regulate ER_GFP__K20 abundance remains to be elucidated.

### The TRAPP complex facilitates the ER-to-Golgi transport of ER_GFP_-K20

The high confident list includes several genes previously implicated in ER-to-Golgi trafficking. These include *TRAPPC1, TRAPPC8*, and *USO1* (Figures 1F, S1E)*. TRAPPC1* and *TRAPPC8* encode subunits of the conserved <u>Tra</u>nsport <u>P</u>rotein <u>P</u>article (TRAPP) complex that is associated with the Golgi (Sacher et al., 1998), while USO1 is a Golgi-localized receptor regulating the inter-cisternal vesicle transport within the Golgi (Sapperstein et al., 1995).

To validate the role of the TRAPP complex in TAQC, we used small hairpin RNAs (shRNAs) to deplete several TRAPP subunits in ER_GFP__K20 cells. Confocal microscopy showed that compared to a scrambled non-targeting shRNA control, shRNAs targeting *TRAPPC1, TRAPPC8, TRAPPC9, TRAPPC10, TRAPPC11* all increased the green fluorescence of ER_GFP__K20 cells (Figures 1G, H). Pulse-chase analysis confirmed that ER_GFP__K20 was stabilized in TRAPPC1-depleted cells (Figures S1F, G). Because depletion of the TRAPP complex did not elevate the level of mCherry in ER_GFP__K20 cells (Figure 1G), we concluded that the TRAPP complex specifically promotes the degradation of translation stalled ER_GFP__K20. These results further corroborate the role of the Golgi system in TAQC.

### SAYSD1 regulates UFM1-dependent degradation of ER_GFP_-K20

Among the high-confidence gene list, a gene named *SAYSD1* (SAYSvFN domain-containing protein 1) encodes an uncharacterized membrane protein (Figure 2A). Like genes in the UFMylation pathway, *SAYSD1* is conserved in metazoan but missing from fungi (Figures S2A). Using siRNA-mediated gene silencing and CRISPR-mediated KO of SAYSD1, we confirmed that depletion of SAYSD1 significantly increased GFP but not mCherry fluorescence in ER_GFP__K20 cells (Figures 2B, C, Figure S2B). Because treating SAYSD1-depleted ER_GFP__K20 cells with Baf A1 caused a much smaller increase in GFP fluorescence than similarly treated control cells (Figure S2C), the ER_GFP__K20 accumulation phenotype in SAYSD1-depleted cells could apparently be attributed to diminished lysosomal degradation. Consistent with this notion, pulse-chase analysis showed that knockdown of SAYSD1 indeed stabilized ER_GFP__K20 (Figure S2D).

**Figure 2.**
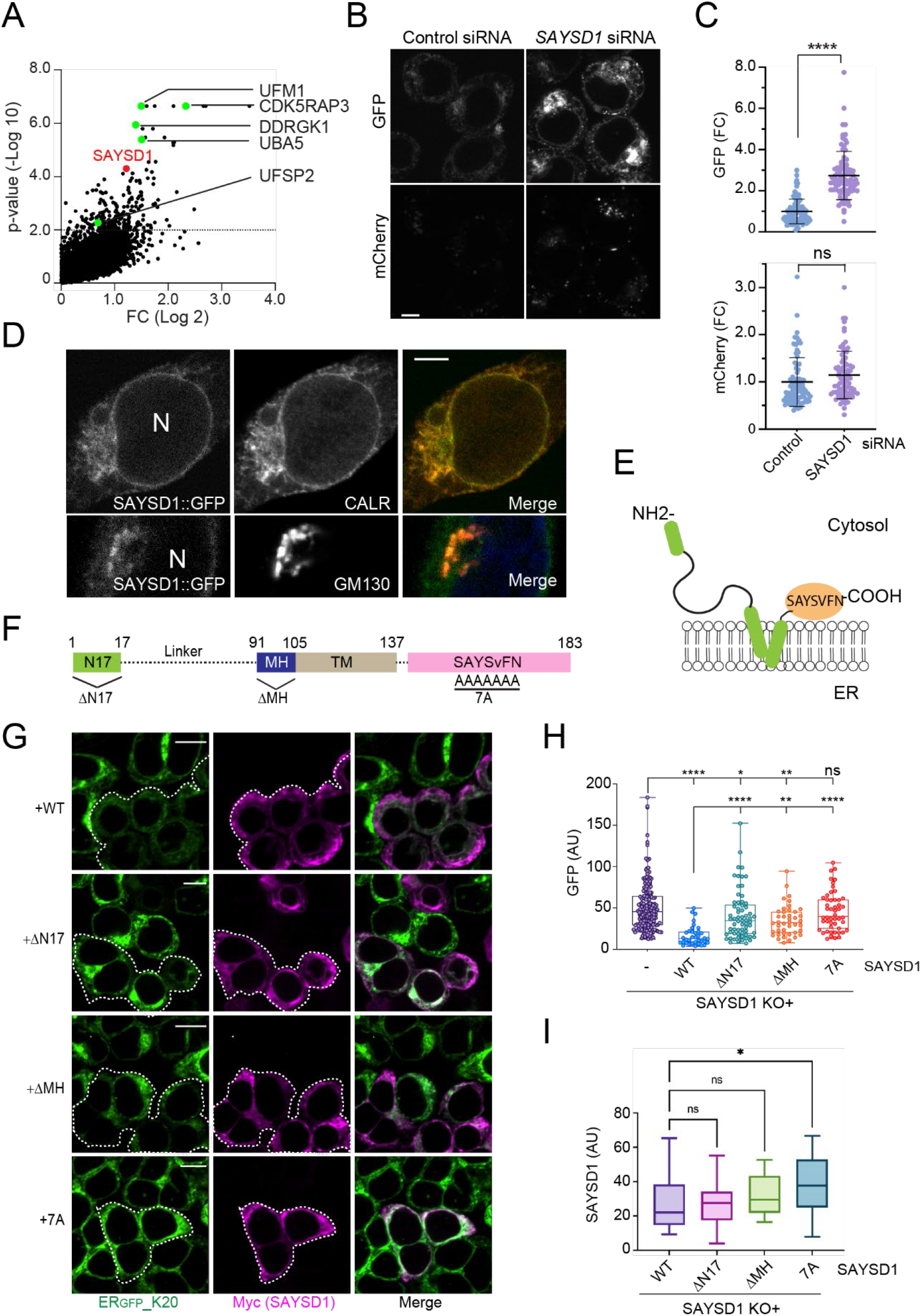
SAYSD1 promotes the degradation of ER_GFP__K20. **(A)** sgRNAs targeting SAYSD1 are enriched in ER_GFP__K20 high cells. The UFM1 pathway genes were highlighted in green. **(B, C)** Knockdown of SAYSD1 increases the ER_GFP__K20 green fluorescence without affecting the full-length read-through product. **(B)** Shown are ER_GFP__K20 stable 293T cells transfected with control or SAYSD1 siRNA for 48 h. Scale bar, 5 μm. The graphs in **(C)** show the quantification of the GFP and mCherry fluorescence in individual cells, respectively. Error bars indicate means ± SD, **** *p*<0.0001 by unpaired Student’s *t*-test; ns, not significant. n=3 independent experiments. **(D)** SAYSD1 is mainly localized to the ER. 293T cells expressing GFP-tagged SAYSD1 from the endogenous locus were fixed and stained with GM130 (bottom panels) or Calreticulin (top panels) antibodies. N, nucleus. Scale bar, 5 μm. **(E)** The predicted membrane topology of SAYSD1. **(F)** The domain structure of SAYSD1 and the mutants used in the rescue study. **(G-I)** Expression of Myc-tagged wildtype (WT) SAYSD1 but not mutants lacking either the SAYS_ν_FN motif (7A), the N-terminal 17 residues (ΔN17), or the middle helical segment (ΔMH) in SAYSD1 knockout (KO) cells restores the degradation of ER_GFP__K20. **(G)** Representative images from the experiments. Dashed lines indicate SAYSD1-positive cells. Scale bars, 10 μm. The graph in **(H)** shows the quantification of GFP fluorescence in individual cells. Error bars indicate means ± SD; **** *p*<0.0001, ** p<0.01, * p<0.05 by one-way Anova. ns, not significant. n= 4 independent experiments. **(I)** Quantification of SAYSD1 expression in **G**.

As expected, neither UFM1 nor SAYSD1 KO affected the steady state expression of ER_GFP__K0, which lacks the ribosome stalling sequence (Figure S2E). However, Baf A1 treatment led to more pronounced accumulation of ER_GFP__K0 in UFM1 and SAYSD1 KO cells than in WT cells, suggesting that defects in UFM1- and SAYSD1-mediated TAQC may induce a compensatory mechanism to degrade damaged ER.

SAYSD1 KO did not affect the expression of either UFM1 or UFMylating enzymes such as UFC1, UFL1, or UFBP1/DDRGK1, nor did it reduce UFMylated ribosome (Figure S2F), suggesting that SAYSD1 may act downstream of ribosome UFMylation. Since UFM1 deficiency has been linked to ER stress induction, we treated cells with ER stress inducers thapsigargin and tunicamycin but found no effect on ER_GFP__K20 (Figure S2G). Together, these results rule out the possibilities that the TAQC defect in SAYSD1-depleted cells is caused by ER stress or lack of ribosome UFMylation.

We next determined the subcellular localization of SAYSD1 using cells bearing a GFP tag at the carboxyl terminus of endogenous SAYSD1 (Figure S2H). Confocal microscopy showed that SAYSD1 is present mostly in the ER, but a fraction was found in perinuclear puncta marked by the Golgi protein GM130 (Figure 2D), suggesting that it may cycle between these membrane compartments. By contrast, ER_GFP__K20 was accumulated mostly in perinuclear puncta in SAYSD1-depleted cells, which were largely co-localized with the ER marker Calreticulin (Figure S2I). These findings implicate SAYSD1 in the export of translation-stalled proteins from the ER during TAQC.

SAYSD1 is predicted to contain a kinked transmembrane domain (TMD) with both the N- and C-termini facing the cytosol (Figure 2E). A short helical segment precedes the TMD, which is followed by a highly conserved SAYSvFN-containing domain (SACD) (Figure 2F, Figure S2A). While the accumulation of ER_GFP__K20 in SAYSD1-depleted cells could be significantly reversed when wildtype (WT) SAYSD1 was re-expressed, expression of the SAYSD1-7A mutant (residues in the SAYSvFN motif mutated to alanine), or mutants lacking either the amino-terminal 17 residues (ΔN17) or the middle helical segment (ΔMH) (Figure 2F) failed to restore ER_GFP__K20 to a level comparable to that in WT SAYSD1-expressing cells (Figures 2G-I). This result suggests a critical role for these conserved motifs in TAQC.

### UFM1-dependent association of SAYSD1 with a stalled nascent chain-ribosome complex

If SAYSD1 directly participates in TAQC, it may interact with the Sec61 translocon. Indeed, immunoprecipitation of Sec61 β, a key component of the ER translocon readily co-precipitated endogenous SAYSD1 but not abundant cytosolic proteins such as p97 (Figure 3A). The interaction of SAYSD1 with the translocon was further supported by co-sedimentation of endogenous SAYSD1 with Sec61β by sucrose gradient centrifugation (Figure S3A). Furthermore, reciprocal immunoprecipitation of GFP-tagged SAYSD1 from *SAYSD1::GFP* cells revealed that SAYSD1 not only interacted with the translocon but also the UFMylation ligase UFL1 (Figure S3B). Interestingly, ribosomes were not significantly co-precipitated with SAYSD1-GFP under these conditions (Figure S3B, lane 2 vs. 1). However, when cells were exposed to anisomycin (ANS), a translation elongation inhibitor that causes ribosome UFMylation (Wang et al., 2020), ribosomes could now be detected in complex with SAYSD1 (Figures S3B, lanes 3, 4; S3C). Upon ANS treatment, we also observed an increased association of SAYSD1 with Sec61β but not UFL1.

**Figure 3.**
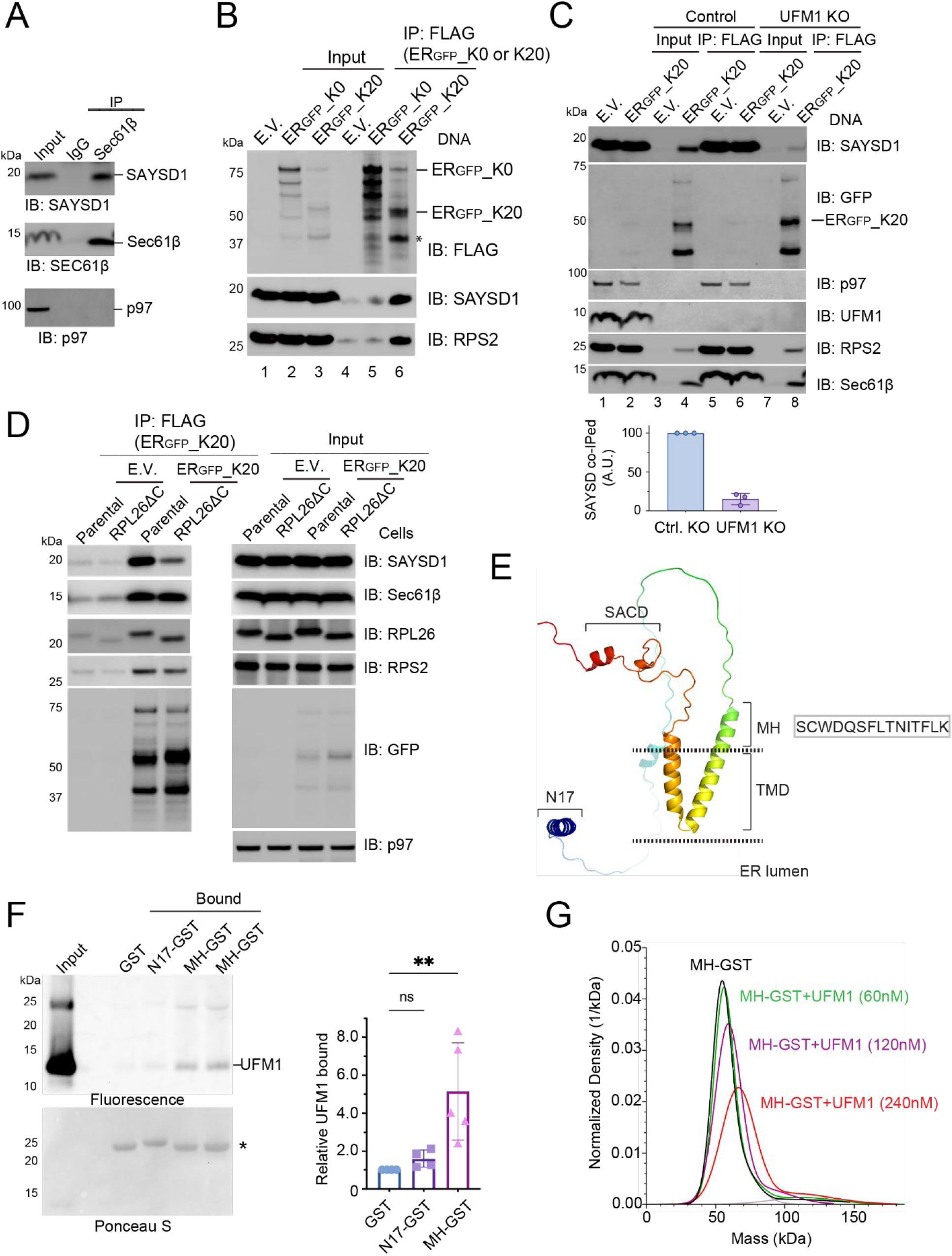
SAYSD1 preferentially engages translocation-stalled nascent chain-ribosome complexes in a UFMylation dependent manner. **(A)** Co-immunoprecipitation of endogenous SAYSD1 with the Sec61 translocon component Sec61β from 293T cells. **(B)** SAYSD1 associates with ER_GFP__K20 in a translation-stalling dependent manner. 293T cells transfected with an empty vector (EV), ER_GFP__K20, or ER_GFP__K0 were subject to immunoprecipitation by FLAG beads. The asterisk indicates the stalled ER_GFP__K20 species. **(C)** The association of SAYSD1 with ER_GFP__K20 is dependent on UFM1. Control or UFM1 CRISPR knockout (KO) cells transfected as indicated were subject to FLAG pulldown and immunoblotting. The graph shows the quantification of three independent experiments. **(D)** The association of SAYSD1 with ER_GFP__K20 requires the RPL26 C-tail. As in **C**, except that cells that have the endogenous RPL26 C-terminal tail deleted by CRISPR-mediated homologous recombination were used. **(E)** The predicted SAYSD1 structure by alphaFold. SACD (<u>SA</u>YSvFN-<u>c</u>ontaining <u>d</u>omain); TMD, transmembrane domain. **(F)** SAYSD1 binds the MH domain of UFM1 directly. The indicated GST-tagged protein or GST immobilized on glutathione beads were incubated with Atto^565^-labeled UFM1. The precipitated proteins were SDS-PAGE fractionated and detected by a fluorescence scanner (top) or Ponceau S staining (bottom). The graph shows the quantification of 4 independent experiments. Error bars, means ± SD; ** p<0.01 by unpaired student t-test. n=4. **(G)** Mass photometry shows a direction interaction of MH-GST with UFM1.

To further probe the engagement of SAYSD1 with the stalled nascent chain-translocon-ribosome complex, we immunoprecipitated ER_GFP_-K20, or as a control, ER_GFP__K0, which lacks the ribosome-stalling sequence but otherwise is identical to ER_GFP__K20 (Wang et al., 2020). As expected, immunoblotting detected a strong interaction of ribosome (as indicated by the ribosome subunit RPS2) with ER_GFP__K20 but not ER_GFP__K0 or empty beads (Figure 3B, lower panels, lane 6 vs. 5, 4). ER_GFP__K20 immunoprecipitation also brought down endogenous SAYSD1 and Sec61β, whereas ER_GFP__K0 only co-precipitated a negligible amount of SAYSD1 (Figures 3B, lane 6, 3C, lane 4). Interestingly, the interaction of SAYSD1 with ER_GFP__K20 was significantly reduced in UFM1 knockout cells (Figure 3C, top panel, lane 4 vs. 8) and in cells lacking the UFMylation site on RPL26 (RPL26 ΔC) (Figure 3D). By contrast, UFM1 depletion or knockout of the RPL26 UFMylation site did not affect the interaction of ER_GFP__K20 with either ribosome or Sec61β (Figures 3C, D, lower panels). Collectively, these results suggest a dynamic interaction between SAYSD1 and the ER translocon, which allows SAYSD1 to engage a stalled nascent chain-ribosome complex upon ribosome UFMylation.

### A middle helical (MH) segment in SAYSD1 binds UFM1 directly

The UFMylation-dependent association of SAYSD1 with ER_GFP_-K20 raised the possibility of SAYSD1 being a UFM1 ‘sensor’. To test this possibility, we expressed and purified GST-tagged proteins containing either the N17, the MH segment, or the C-terminal SACD (138-183) from *E. coli* (Figure S3D). We chose these regions because of sequence conservation and also because they are required for efficient ER_GFP_-K20 turnover (Figure 2). Additionally, alphaFold predicts that these segments are folded independently in the cytosol (Figure 3E) (Jumper et al., 2021). We incubated purified proteins and a GST control with recombinant UFM1. Glutathione bead pulldown showed that only MH-GST consistently pulled down UFM1 whereas GST, N17-GST or GST-SAYSD1137-183 failed to interact with UFM1 (Figures 3F, S3E).

We further confirmed the interaction between MH-GST and UFM1 using single molecule mass photometry. We first examined the effect of UFM1 on the mass of MH-GST using N17-GST as a negative control. As expected, incubating UFM1 with N17-GST did not change the mass of the latter (Figure S3F). By contrast, incubating UFM1 with MH-GST resulted in a concentration dependent increase in the mass of MH-GST (Figures 3G, S3F), suggesting the formation of a complex consisting of UFM1 and MH-GST. The specificity of the interaction was further supported by the observation that MH-GST did not bind ubiquitin under the same condition (Figure S3F). Collectively, these results demonstrate a direct and specific interaction between the MH segment of SAYSD1 and UFM1, which provides a molecular basis of UFM1-dependent engagement of SAYSD1 with the stalled nascent chain-ribosome complex.

### A conserved SAYSD1 N-domain binds ribosome to promote TAQC

The N17 domain of SAYSD1 is predicted to form a helix with a conserved phenylalanine (F) followed by four positively charged residues (Figures 4A, B). Interestingly, the bacterial trigger factor uses a similar N-terminal ‘signature motif’ to interact with ribosome (Figure 4A) (Kramer et al., 2002). Likewise, the ribosome binding site in nascent polypeptide-associated complex (NAC) is formed by a stretch of positively charged residues (RRKxxKK) at the N-terminus (Wegrzyn et al., 2006). These observations suggest that the N17 motif of SAYSD1 may also bind ribosome. To test this idea, we performed GST pulldown after incubating recombinant GST-N17 or GST (control) with a 293T cell extract. Immunoblotting showed that GST-N17 but not GST readily coprecipitated with ribosomes (Figure 4C). Several additional lines of evidence further demonstrate N17 as a ribosome binding domain that recognizes ribosome independent of UFMylation. First, although immunoblotting with UFM1 antibodies showed co-precipitation of N17 with UFM1-conjugated RPL26, immunoblotting with RPL26 antibodies also detect unmodified RPL26 in association with GST-N17 (Figure 4C, top panel). Second, the association of GST-N17 with ribosome was not affected by UFM1 depletion or by treating cells with anisomycin, which abolished and increased UFMylated ribosome, respectively (Figures S4A, B). Likewise, knockout out of the ribosome UFMylation site in RPL26 had no effect on this interaction (Figure S4C). Importantly, ribosome purified from rabbit reticulocyte lysate similarly interacted with GST-N17 (Figure 4D), suggesting that N17 interacts directly with ribosomes. This interaction is highly specific because GST-N17 did not pull down the abundant p97 ATPase or HSP90 (Figures 4C, E). Importantly, SAYSD1 variants carrying mutations on conserved residues (3A: L5A, F8A, R12A or E7A_R14A) lost the ribosome binding activity completely (Figure 4E). These results suggest that N17 may function as a ribosome binding site during TAQC.

**Figure 4.**
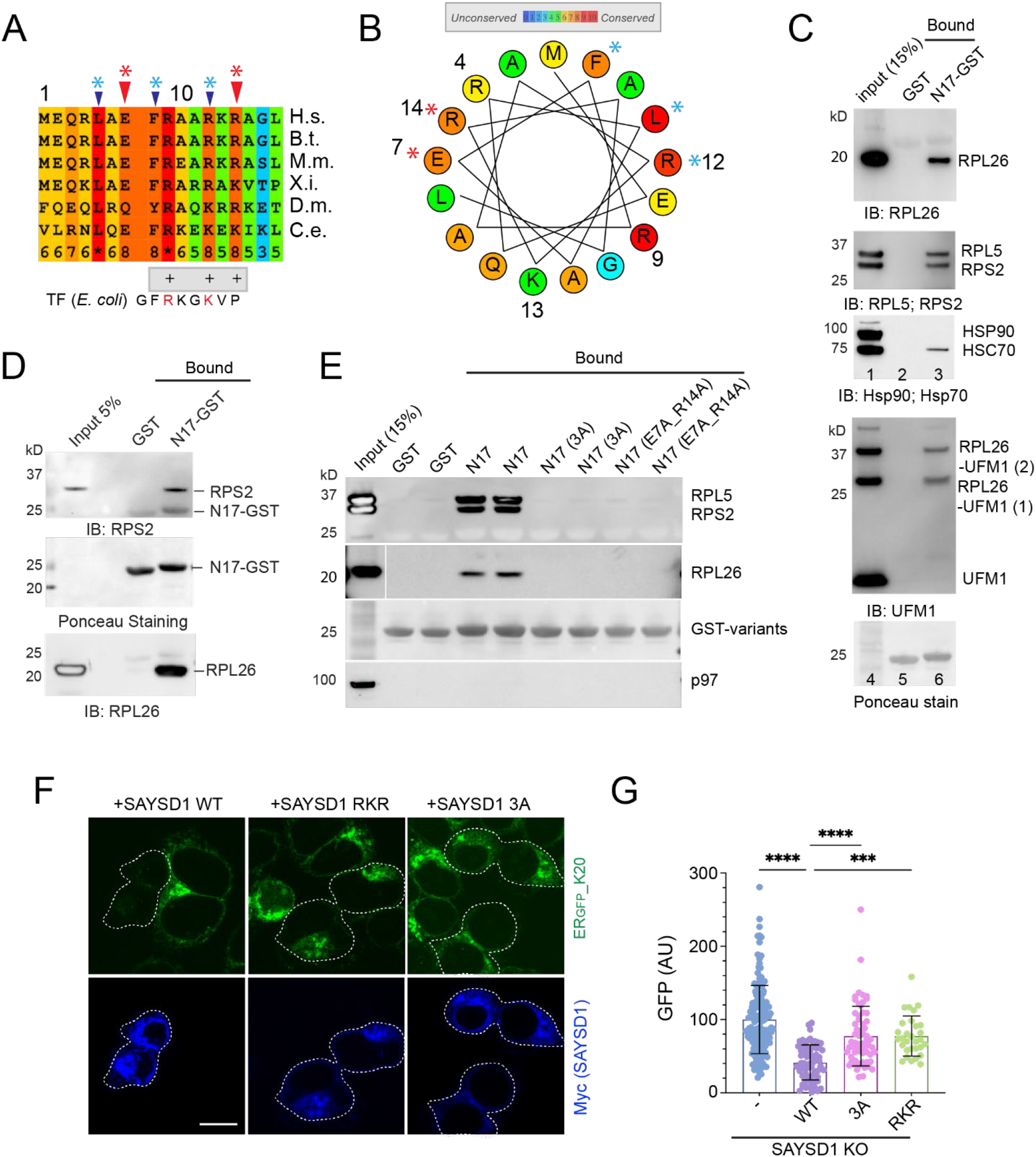
SAYSD1 recognizes ribosome directly via an N-terminal segment. **(A)** Sequence alignment of the N17 domain of SAYSD1. The arrows indicate the two groups of conserved residues mutated in the study. **(B)** A helix wheel view of the SAYSD1 N17 domain. Colors indicate the degree of conservation. Asterisks indicate residues mutated. **(C, D)** GST-N17 interacts with ribosome directly. **(C)** GST-N17 or GST immobilized on glutathione beads were incubated with 293T cell extract. Precipitated proteins were analyzed by immunoblotting with the indicated antibodies. **(D)** As in **C**, except that ribosome purified from rabbit reticulocyte lysate was used. **(E)** The conserved residues in N17 are required for ribosome binding. The indicated GST-tagged N17 variants or GST was immobilized in duplicate and incubated with 293T cell lysate. The precipitated proteins were analyzed by immunoblotting using indicated antibodies. **(F, G)** SAYSD1 ribosome binding is required for efficient turnover of ER_GFP__K20. (**F**) SAYSD1 KO cells expressing ER_GFP__K20 were transfected with the indicated DNA and imaged. SAYSD1-positive cells are highlighted by dashed lines. Scale bar, 10 μm. The graph in (**G**) shows the quantification of randomly imaged cells from two independent experiments. Error bars, means ± SD; **** *p*<0.0001; *** p<0.001 by unpaired Student’s *t*-test.

To confirm the role of ribosome binding in TAQC, we expressed the SAYSD1 3A and another mutant that had three positively charged residues (RKR) mutated to alanine in ERGFP_K20-expressing SAYSD1 KO cells. Compared to WT SAYSD1, these mutants were significantly less effective in restoring the degradation of ER_GFP__K20 (Figures 4F, G). Thus, ribosome binding by the N17 domain plays a critical role in TAQC.

### UFM1 and SAYSD1 eliminate translation-stalled collagen I in mammalian cells

To elucidate the physiological function of TAQC, we investigated the role of UFM1 and SAYSD1 in collagen biogenesis. Collagens are the most abundant secretory proteins in metazoan (Gelse et al., 2003). Most collagen molecules have a triple-helical domain (THD) that contains many repeated “PPG” motifs whose translation is known to cause ribosome pausing/stalling (Huter et al., 2017; Peil et al., 2013). This unique feature renders collagens a good candidate as an endogenous TAQC substrate.

To test whether the translation of the THD of collagens could cause translation arrest, we cloned the human collagen A1 (ColA1) THD-coding sequence between a GFP- and a mCherry-coding sequence (Figure 5A). A viral 2A sequence was inserted both upstream and downstream of the THD sequence. This sequence causes ribosomes to re-initiate translation, resulting in two equally translated polypeptides from a single mRNA (Ryan et al., 1991). In our design, the translation products should include GFP, Col1A1 THD, and mCherry, all in equal amount if the Col1A1 THD sequence did not stall the ribosome. We used the ribosome stalling poly-adenine sequence (K20) as a positive control and a sequence with no stalling motif (K0) as the negative reference. Flow cytometry analyses of cells expressing these reporters showed that the poly-A sequence reduced the mCherry/GFP ratio by about 75% when compared to cells over-expressing GFP-K0, consistent with K20 being a strong stalling motif (Juszkiewicz and Hegde, 2017). The Col1A1 THD sequence reduced the mCherry/GFP ratio by about 30% (Figure 5B), suggesting that it does cause translation arrest, albeit less efficiently than the poly-A sequence.

**Figure 5.**
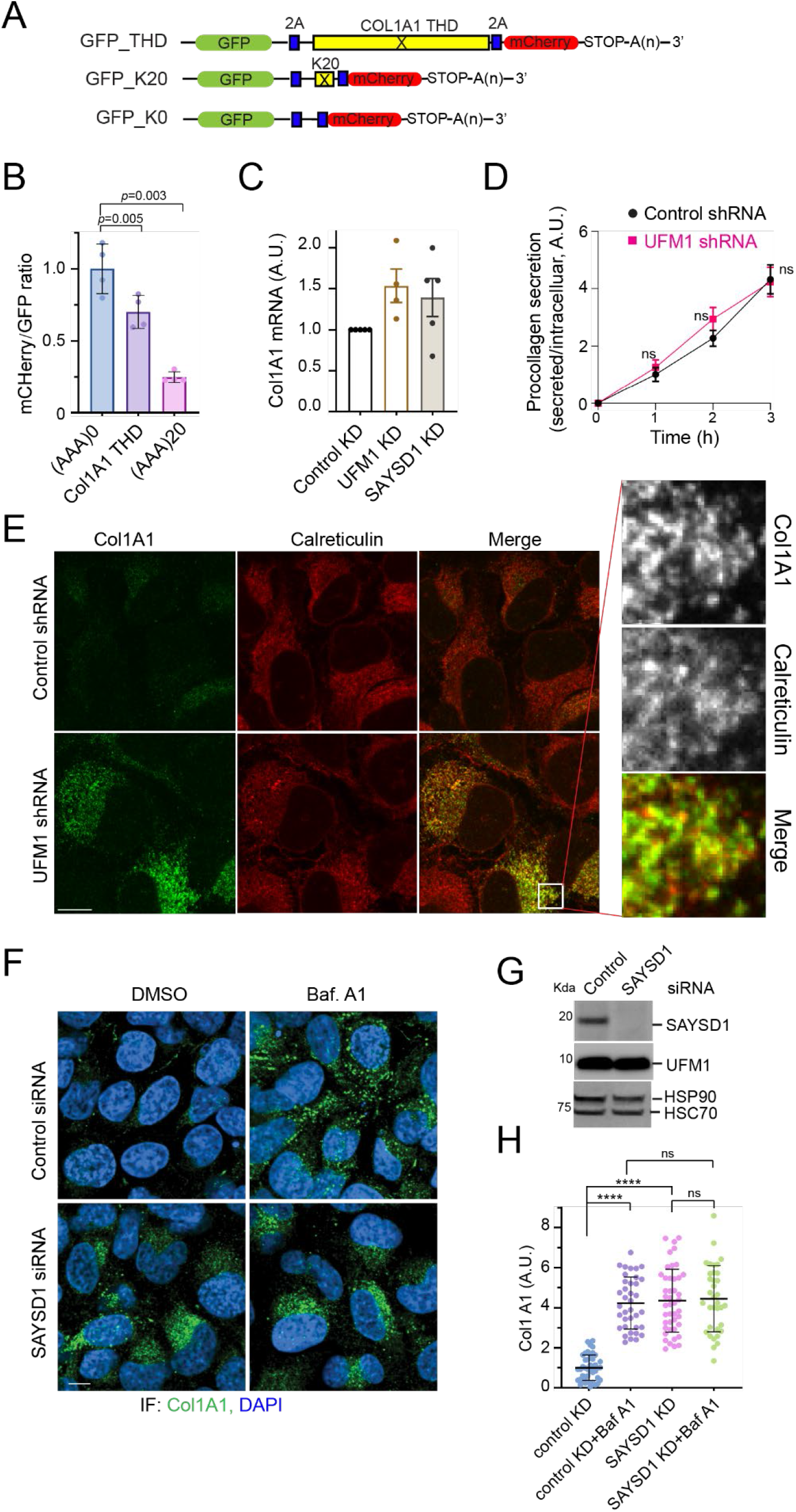
UFM1- and SAYSD1-dependent TAQC degrades translation-stalled Col1A1. **(A)** A schematic view of the translation stalling reporters and the K0 control. **(B)** The translation of the Col1A1 triple helical domain (THD) causes ribosome stalling. 293T cells were transfected with the indicated reporters. The ratio of mCherry vs. GFP was determined by flow cytometry. Error bars, means ± SD; p-values are determiend by one-way ANOVA followed by Dunnett’s multiple comparisons test; n=4 independent experiments. **(C)** UFM1 or SAYSD1 knockdown does not significantly increase Col1A1 mRNA as determined by qRT-PCR. n=3 independent experiments. **(D)** UFM1 depletion does not affect Col1A1 secretion. Conditioned medium from control or UFM1-depleted U2OS cells were analyzed by ELISA. Error bars indicate means ± SD. ns, not significant by unpaired student t-test, n=3 independent experiments. **(E)** UFM1 depletion stabilizes Col1A1 in the ER. Control and UFM1 knockdown U2OS cells were stained by Col1A1 (green) and Calreticulin (red) antibodies. Scale bar, 10 μm. Right panels show an enlarged view of the boxed area. **(F-H)** Baf A1 treatment does not further enhance SAYSD1-depletion induced Col1A1 accumulation. **(F)** Control and SAYSD1 knockdown U2OS cells were treated with DMSO as a control or with Baf A1 (200 nM, 5 h), and stained by Col1A1 antibodies (Green) or DAPI (Blue). Scale bar, 10 μm. **(G)** A fraction of the siRNA-treated cells in **(F)** were analyzed by immunoblotting to confirm the knockdown of SAYSD1. **(H)** Quantification of the Col1A1 levels in **F**. Error bars, means ± SD; **** p<0.0001, by one-way ANOVA with Dunnett’s multiple comparisons test. n=3 independent experiments.

We next analyzed the level of endogenous Col1A1 in control and *UFM1* knockdown human osteosarcoma U2OS cells, which naturally produce and secrete collagen 1 (Forrester et al., 2019). Immunostaining showed that *UFM1* knockdown increased Col1A1 protein level by ~5-fold (Figures S5A, B), which was at least partially regulated at a post-transcriptional level because qRT-PCR revealed only a small increase in the Col1A1 mRNA level by UFM1 depletion (Figure 5C). The accumulation of Col1A1 was not caused by lack of protein secretion because ELISA analysis of conditioned medium detected no difference in secreted Col1A1 between control and *UFM1* knockdown cells (Figure 5D). Similar to UFM1 depletion, Baf A1 treatment also caused Col1A1 to accumulate; Combining Baf A1 treatment with *UFM1* knockdown did not lead to further Col1A1 accumulation compared to Baf A1 treatment or UFM1 depletion alone (Figures S5B, C). These results suggest that Col1A1 accumulation in UFM1-depleted cells is mainly caused by reduced lysosomal degradation. Importantly, re-expressing UFM1 in *UFM1* knockdown cells drastically reduced Col1A1 levels (Figures S5D, E), demonstrating a specific causal link between UFM1 depletion and Col1A1 stabilization.

Dual-color fluorescence confocal analyses showed that Col1A1 was mainly co-localized with the ER protein Calreticulin in *UFM1* knockdown cells (Figure 5E). By contrast, in Baf A1-treated cells, Col1A1 accumulated in vesicle-like puncta, which were mostly free of Calreticulin (Figure S5F). These results are consistent with the established role of UFM1 in TAQC, suggesting that UFMylation also facilitates the export of stalled Col1A1 for lysosomal degradation.

Like in UFM1-depleted cells, knockdown of *SAYSD1* also caused a significant accumulation of Col1A1 protein (Figures 5F-H), but no significant increase in its mRNA (Figure 5C). Baf A1 treatment did not further enhance Col1A1 accumulation in *SAYSD1* knockdown cells (Figures 5F-H). Altogether, these results indicate that UFM1- and SAYSD1-mediated TAQC constitutively targets a fraction of newly synthesized Col1A1 for lysosomal degradation (see discussion).

### UFMylation and SAYSD1 promote collagen biogenesis in flies

To study the *in vivo* function of TAQC, we first performed immunostaining using a UFM1 specific antibody to determine the expression of UFM1 in fat body (FB) and salivary gland (SG) of *Drosophila* third instar larvae. These tissues are dedicated to protein secretion with UFM1 readily detectable by confocal microscopy (Figure S6A). Interestingly, we observed low UFM1 staining in immature SG, but high in mature, secretion-competent SG bearing a sizable lumen (Figure S6A, iii vs. ii and i). The observation suggests a conserved role for UFMylation in ER protein biogenesis in flies.

To further study TAQC in flies, we generated transgenic flies carrying the *ER_GFP__K20* reporter downstream of an UAS regulatory element. *ER_GFP__K20* flies were crossed to a *GMR-Gal4* enhancer line, which specifically expressed the UAS-binding transcription factor Gal4 in photoreceptor cells of larval eye discs. This resulted in the expression of ER_GFP__K20 in a similar pattern as Gal4. Similar as in mammalian cells, we detected weak GFP and no mCherry in eye discs of *GMR>ER_GFP__K20* flies (Figures 6A, panel i, 6B), suggesting that translation stalling by the poly-A segment also occurs in flies. Knockdown of *UFM1* significantly increased the GFP signal with no impact on mCherry, so did depletion of *TRAPPC11* (Figures 6A, panels ii, iii vs. i, 6B). Similar results were obtained when we used the *Cg-Gal4* line, in which the Gal4 expression is driven by a FB specific promoter of the *COL4* gene (*Viking*) (Figures S6B, C). Collectively, these results suggest that UFM1-dependent TAQC is conserved in *Drosophila*.

**Figure 6.**
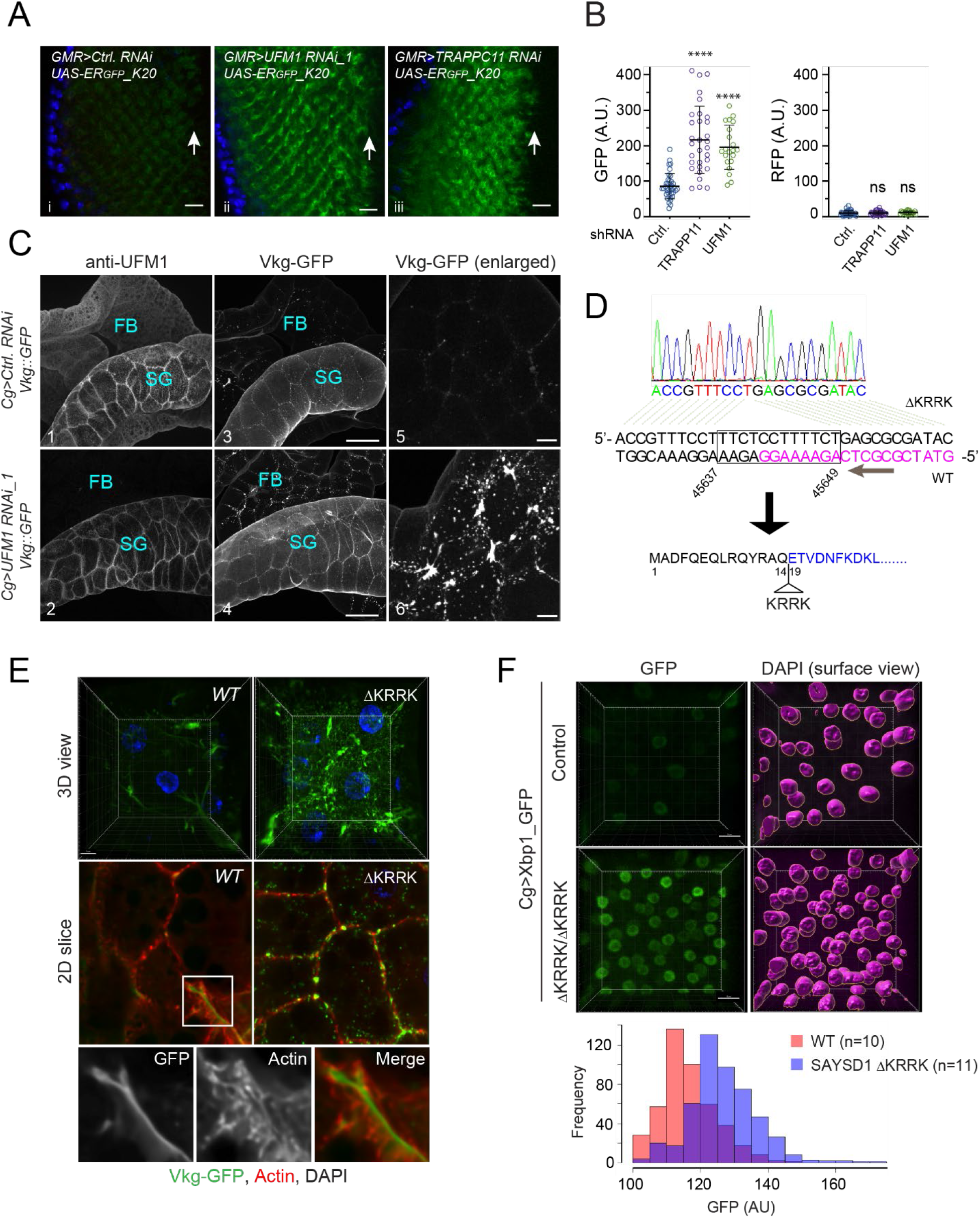
UFMylation and SAYSD1 regulate TAQC and ER protein biogenesis in *Drosophila*. **(A)** Knockdown of *UFM1* or *TRAPPC11* in photoreceptor cells caused accumulation of ER_GFP__K20. Scale bars, 10 μm. **(B)** Quantification of the fluorescence intensity in individual eye discs as shown in **A**. Error bars indicate means ± SD. **** *p*<0.001 by one-way ANOVA with Dunnett’s multiple comparisons test. ns, not significant, n=4 independent experiments. **(C)** Fat body (FB) specific knockdown of UFM1 causes Viking-GFP (Vkg-GFP) to accumulate in FB. Salivary gland (SG)-associated FB from larvae of the indicated genotypes were stained with UFM1 antibodies (panels 1, 2). Shown are maximum projected views of the confocal sections. Panels 5, 6 show an enlarged view of Viking-GFP in FB. Scale bars: panel 1-4, 100 μm; panel 5-6, 20 μm. **(D)** DNA sequencing to validate the genotype of SAYSD1 CRISPR flies. DNA from homozygous SAYSD1 CRISPR 2^nd^ instar larvae was used to amplify SAYSD1 genomic locus, which was then sequenced. Magenta indicates the sgRNA sequence. **(E)** As in **C**, except that FB from second instar larvae of wildtype (WT) and SAYSD1 homozygous ΔKRRK mutants were stained with an actin dye (red) were imaged. All flies also bear the Vkg::GFP reporter. Scale bar, 10 μm. **(F)** ER stress is induced in SAYSD1 ΔKRRK homozygous flies. FB from control or SAYSD1 ΔKRRK second instar larvae bearing an ER stress reporter (Xbp1-GFP) were stained with DAPI and imaged by 3D confocal microscopy. 3D surfaces were rendered by Imaris for individual nucleus, which were used to quantify the GFP signal. The graph shows the quantification results from two independent experiments. Scale bar, 15 μm.

To test whether TAQC regulates collagen biogenesis in flies, we knocked down *UFM1* in COL4-producing FB in larvae that had a GFP-encoding sequence inserted in the endogenous *Viking* locus (Viking-GFP) (Morin et al., 2001). *Drosophila* COL4 consists of two polypeptide chains, COL4A1 and COL4A2 (also named Viking), which are a major component of the basement membrane (Yurchenco and Furthmayr, 1984). COL4 is secreted from FB and then incorporated into the basement membranes of imaginal discs and guts to maintain tissue homeostasis (Pastor-Pareja and Xu, 2011). To rule out potential off-target effects, we used two UFM1 shRNA lines. As a negative control, flies carrying mCherry shRNA were used. Immunostaining with UFM1 antibodies confirmed that UFM1 was specifically depleted in FB of Cg>UFM1 RNAi flies but its level in nearby tissues such as SG remained unchanged (Figure 6C, panel 2 vs. 1). Confocal microscopy detected only weak Viking-GFP signal in fat bodies in control larvae (panels 3, 5), suggesting that newly synthesized COL4, if not secreted, is degraded rapidly. By contrast, we observed many Viking-GFP-containing puncta in UFM1-depleted fat bodies (Figures 6C, panels 4, 6 vs. 3, 5; S6D). This phenotype could not be attributed to a secretion defect because the Viking-GFP signal in the basement membranes around SG and guts (see below) was comparable between control larvae and larvae with FB specific UFM1 depletion (Figure 6C). Thus, a significant fraction of COL4 appears to be downregulated in a UFM1 dependent manner during *Drosophila* development.

To validate the role SAYSD1 in collagen biogenesis in flies, we used CRISPR/Cas9 to generate *SAYSD1* loss-of-function mutant flies (see method). An initial screening of more than 30 individual CRISPR lines identified a line that harbors a 12-nucleotide deletion in the *SAYSD1* locus. The resulting allele encodes a protein missing four charged residues (ΔKRRK) in the ribosome binding N17 domain (Figure 6D). Although homozygous ΔKRRK flies died before the third instar larval stage, Viking-GFP accumulation was observed by 3D confocal microscopy in FB of second instar *SAYSD1 ΔKRRK* mutant larvae. In wildtype FB, Viking-GFP was detected as a weak smooth signal in spaces between FB cells, suggesting that it is mostly secreted. In SAYSD1 ΔKRRK homozygous FBs, Viking-GFP was detected as bright green puncta both in and outside the cell (Figure 6E). Thus, like UFM1, SAYSD1 also contributes to TAQC of COL4 in flies.

To further evaluate the role of SAYSD1 in ER protein biogenesis, we crossed Cg>Xbp1-GFP flies to either control or SAYSD1 ΔKRRK flies. The Xbp1-GFp reporter contains an unconventional intron from Xbp1. Only when this intron is spliced out during ER stress, will the downstream GFP be translated in frame with Xbp1. Thus, the level of GFP expression reflects the level of ER stress. As anticipated, FB cells from SAYSD1 ΔKRRK larvae have a much higher GFP expression compared to control WT FB (Figure 6F). Because ER stress upregulation was reported in UFM1 knockout cells (Cai et al., 2015; Lemaire et al., 2011; Zhang et al., 2012), these results further corroborate the notion that UFM1 and SAYSD1 function together to promote ER protein homeostasis.

### TAQC controls the quality of secreted collagen in flies

Since deficiency in TAQC does not reduce collagen secretion, but leads to reduced lysosomal degradation, we postulated that translation-stalled Viking-GFP, if not efficiently degraded, may escape from UFM1 or SAYSD1 deficient FB. Incorporation of such defective collagens into the basement membranes might disrupt the basement membrane integrity. To test this idea, we examined the Viking-GFP-containing collagen fibrils on the gut of third instar larvae by confocal microscopy, focusing the proventriculus and middle midgut where Viking-GFP forms stereotyped patterns at this developmental stage (Figure S7A). On the proventriculus of control larvae, Viking-GFP forms six evenly spaced rings, but in larvae that had UFM1 depleted from FB, most proventriculus only have 4 Viking-GFP positive rings, which were often irregularly spaced (Figure 7A, panels 1-3). On the middle midgut, WT larvae have long parallel and evenly spaced longitudinal collagen fibrils (LCFs), which are intersected perpendicularly by regularly spaced circular collagen fibrils (CCFs) (Figures 7A, panels 4, 7, S7A). By contrast, in larvae with FB specific depletion of UFM1, LCFs are often disrupted or formed intersections, while CCFs are curled, irregularly spaced or missing in some areas (Figure 7A, panels 5, 6, 8, 9). These phenotypes were even more prominent in the gut of adult *Cg-Gal4; Viking-GFP/UAS-UFM1 RNAi* flies (Figures S7B, C). As a result, the visceral muscle cells, which are normally attached to Viking-GFP-containing COL4 fibrils, are no longer positioned properly (Figure S7B).

**Figure 7.**
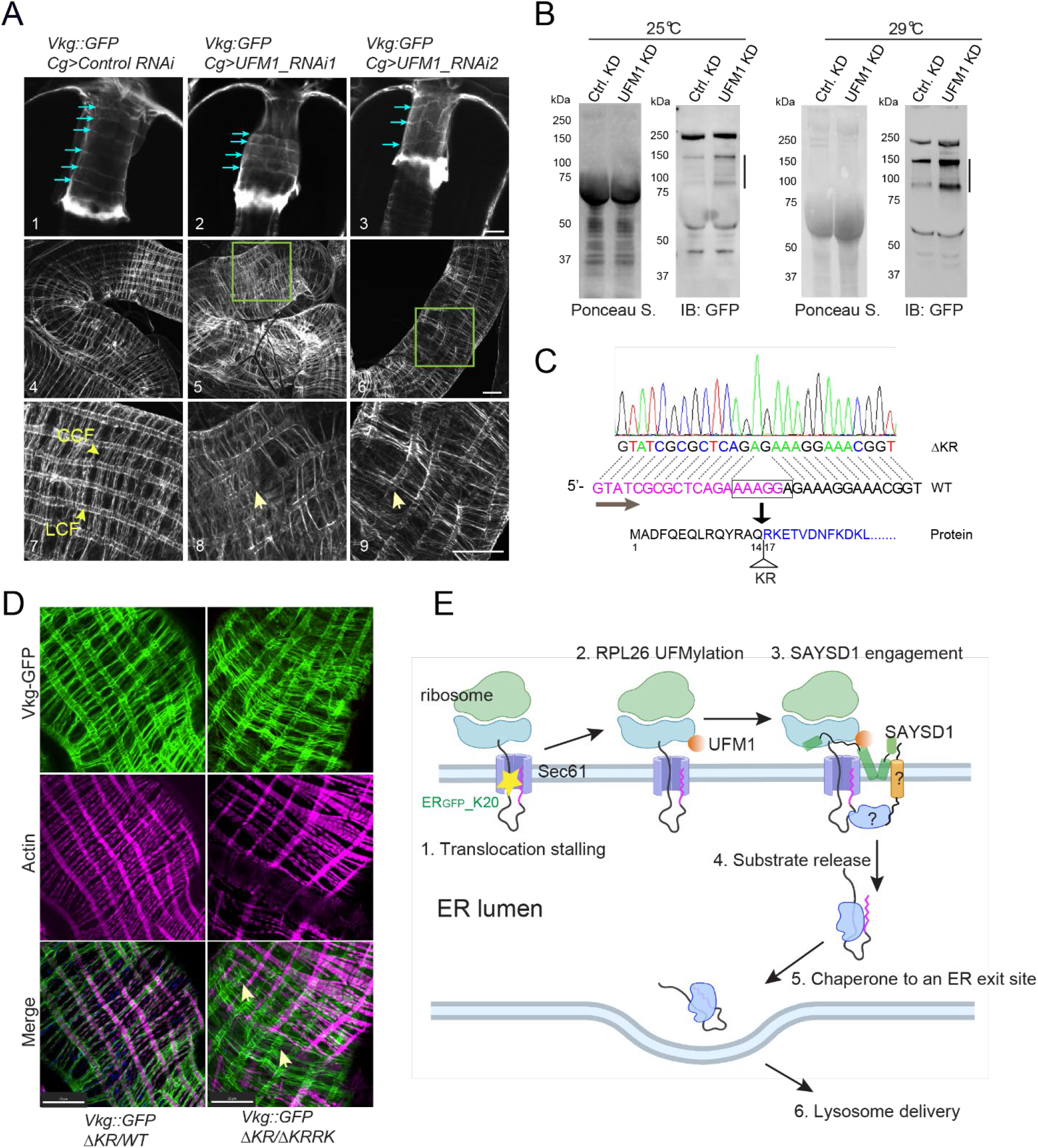
TAQC safeguards collagen biogenesis in *Drosophila*. **(A)** Abnormal Viking deposition by UFM1-depleted FB in fly larvae. Shown are representative confocal images of Vkg-GFP on proventriculus (Panels 1-3) or middle midgut (Panels 4-9) from third instar larvae of the indicated genotypes. Blue arrows in panels 1-3 indicate collagen fibrils around the proventriculus. Arrows in panels 8, 9 indicate disrupted LCFs. Panels 7-9 are enlarged views. Scale bars, 20 μm. **(B)** Defective Vkg-GFP accumulation in hemolymph in larvae with FB specific UFM1 depletion. The lines indicate truncated Viking-GFP species. **(C)** Identification of a second SAYSD1 allele deleting two charged residues in the N17 domain. **(D)** Abnormal Viking-containing basement membranes on middle midgut of fly larvae bearing the indicated mutant SAYSD1 alleles. The guts from third instar larvae were stained by phalloidin to label actin (magenta). Arrows indicate Vkg-GFP-containing fibrils that are detached from muscle cells. Scale bars, 50 μm. **(E)** A model of ribosome UFMylation sensing by SAYSD1 in TAQC. Created by Biorender.com

To further prove that defective collagens are indeed secreted from UFM1 deficient FB, we used immunoblotting to examine the secreted Viking-GFP in hemolymph collected from either control or FB specific UFM1 knockdown larvae. Consistent with our hypothesis, we detected several truncated Viking-GFP species that were elevated by FB specific UFM1 KD (Figure 7B). By contrast, the secretion of full-length Viking was largely unaffected.

To examine the role of SAYSD1 in basement membrane quality control, we screened additional CRISPR lines to identify a mutant that has only two positively charged residues (ΔKR) deleted from the ribosome-binding N17 domain (Figure 7C). Flies bearing homozygous ΔKR alleles or trans-heterozygous ΔKR and ΔKRRK alleles are viable, probably because the smaller deletion had a less severe impact on the SAYSD1 function. Nonetheless, trans-heterozygous SAYSD1 third instar larvae (ΔKR/ΔKRRK) have pronounced defects in collagen quality control and basement membrane assembly. These include Viking-GFP accumulation in FB cells and irregularly positioned Viking-GFP fibrils with many detached from the underlying muscle cells along middle midgut (Figure 7D). Altogether, these results establish a critical role for UFM1- and SYASD1-mediated TAQC in the elimination of defective collagen molecules, which maintains the structural integrity of the basement membrane in *Drosophila*.

Intriguingly, despite the abnormal basement membrane morphology, flies with FB specific UFM1 knockdown developed normally with no discernable phenotype upon eclosion. FB specific UFM1 knockdown also did not affect the fly life span under normal growth conditions. However, when raised at an elevated temperature (29 °C), flies with FB specific UFM1 knockdown had a shortened life span (Figure S7D vs. E). These findings suggest that the basement membrane integrity is not essential for fly development but confers stress resistance in adult flies.

## Discussion

In this study, we used an unbiased genome-wide screen to identify genes involved in TAQC-mediated elimination of polypeptides stalled during co-translational protein translocation. Our screen re-discovers the UFMylation pathway, underscoring the significance of this post-translational modification in TAQC. Additionally, we have uncovered an uncharacterized gene named *SAYSD1* that functions together with the UFMylation system in this process. *SAYSD1* encodes a membrane protein, which associates with the Sec61 translocon in a stalling regulated manner. It contains a highly conserved TMD, which may be the site of Sec61 binding. This TMD is predicted to form a kinked configuration, allowing a conserved N-terminal segment (N17), a middle helix (MH) before the TMD, and the highly conserved SAYSvFN-containing domain downstream of the TMD to be all exposed in the cytosol. We demonstrate that N17 is a novel ribosome binding domain, and the MH segment forms a UFM1 binding site. Since the UFMylation site on RPL26 is close to the Sec61 translocon (Voorhees et al., 2014), the bipartite mode of ribosome recognition may allow N17 and MH to jointly sense UFMylated ribosomes that are docked on clogged Sec61 translocon (Figure 7E). Although the N17 domain of SAYSD1 can readily interact with ribosome in cell extracts, we did not detect significant association of endogenous SAYSD1 with ribosome in the absence of translocation stalling. Thus, it is possible that this domain is kept in an inactivated form until translocation stalling induces ribosome UFMylation.

Our findings suggest that in TAQC, translocation-stalled polypeptides are released into the ER lumen and then transported to lysosomes via the Golgi. Our screen identifies the TRAPP complex as a facilitator of this process. The TRAPP complex is known to tether ER-derived vesicles to the Golgi apparatus in the canonical ER-to-Golgi trafficking pathway (Barrowman et al., 2010). However, our screen does not recover any COPII components. This may be due to genetic redundancy or unsaturated screen design.

While our findings suggest SAYSD1 as a sensor that monitor the UFMylation status on ribosomes during co-translational protein translocation, molecular details downstream of the SAYSD1-UFM1 interaction are unclear. Intriguingly, TAQC substrates are released from UFMylated ribosomes in a UFM1 independent manner. Nevertheless, their delivery to lysosomes still requires UFM1 and SAYSD1. Thus, ribosome UFMylation might elicit an ‘outside-in’ signal, which allows the ‘sorting’ of faulty substrates away from properly folded proteins that are destined to the cell surface or the cell exterior (Figure 7E). SAYSD1 may interact with a membrane protein with a luminal domain, which in turn recruits a downstream chaperone to escort TAQC substrates to the correct destination. Alternatively, the stalled translocon may adopt a special conformation to engage a luminal protein. The exact nature of this signal and how it is deciphered in the ER lumen remain to be elucidated.

Previous studies suggested that a significant fraction (10-20%) of the newly synthesized collagens are rapidly degraded (Bienkowski, 1989). Accordingly, numerous quality control mechanisms such as ERAD and ERphagy have been proposed for elimination of misfolded or aggregated collagens at post-translocation stages (Ito and Nagata, 2021). Our study shows that translocation pausing due to the enrichment of the stalling-prone PPX motifs in collagens is another source of defective collagens endogenously produced in animals. Our data suggest that the delicate mechanism of collagen biogenesis evolved by nature creates an unavoidable translation hardship, necessitating the UFM1- and SAYSD1-dependent TAQC to safeguard collagen biogenesis. This notion would explain the absence of the UFMylation system and SAYSD1 in yeast, which has no orthologous collagen genes. Intriguingly, UFM1 was recently identified as a positive regulator of ERphagy by a CRISPR/Cas9-based genetic screen, and an ERphagy receptor FAM134 was reported to play a role in collagen quality control at the ER in cultured cells (Forrester et al., 2019; Liang et al., 2020; Reggio et al., 2021). These observations raise the possibility that UFMylation-regulated ER cargoes may use both the canonical ER-to-Golgi pathway and ERphagy to reach lysosomes depending on whether an ERphagy receptor is present on the cargo-bearing vesicles. Given the previously established links between collagen and CNS disorders (Gregorio et al., 2018; Palacios-Macapagal et al., 2021), out study hints at the possibility that defective basement membrane assembly and/or abnormal collagen deposition may contribute to neuronal dysfunctions that are associated with genetic lesions in the UFMylation system.

## Limitations of this study

While our results suggest that SAYSD1 acts as a UFM1 sensor to regulate TAQC, we acknowledge that our study has several limitations. First, while we identified a region in SAYSD1 that is sufficient to bind UFM1, we cannot exclude the involvement of other SAYSD1 sequences or yet-to-be identified SAYSD1-binding proteins in this recognition. For tested TAQC reporter, knockout of SAYSD1 only results in partial stabilization of the substrate, suggesting that additional mechanisms may also contribute to TAQC. Additionally, the reported lysosomal degradation mechanism is demonstrated mostly with one model substrate. It is thus possible that other translation-stalled substrates may use other mechanisms (e.g. ERAD or ERphagy) for degradation upon being released into the ER lumen, particularly if they bear additional folding defects or other unstable structural signatures. Along the same line, although our study establishes collagens as the first endogenous substrate whose biogenesis is linked to ribosomes stalling and regulated by UFM1 and SAYSD1, whether translation stalled-collagens use the exact same ER-to-Golgi route as the ER_GFP__K20 reporter for trafficking to lysosomes awaits further characterization.

## STAR Methods

### Key resources table

#### Resource availability

##### Lead contact

Further information and requests for reagents may be directed to and will be fulfilled by lead contact Yihong Ye

##### Materials availability

All unique/stable reagents generated in this study are available from the lead contact with a completed Materials Transfer Agreement.

##### Data and code availability

- This paper does not report original code.
- Any additional information required to reanalyze the data reported in this paper is available from the lead contact upon request.

#### Experimental model and subject details

The 293T, 293FT, U2OS cells were purchased from ATCC. These cells were maintained in Dulbecco’s Modified Eagle Medium (DMEM, Corning) containing 10% fetal bovine serum (FBS) and antibiotics (penicillin/streptomycin, 10 U/mL) at 37 °C in a 5% CO2 humidified atmosphere.

Fly lines were ordered from the Bloomington Drosophila Stock Center. UAS-ER_GFP__K20 transgenic flies were generated using a strain (R8622) carrying an attB integration site at 68A4 by Rainbow Transgenic. Flies were maintained on standard food in a 25 °C incubator.

#### Method details

##### Cell culture and transfections

To facilitate procollagen biogenesis in U2OS cells, 50 μg/mL L-ascorbic acid was added in the culture medium. Plasmid transfection was performed using the TransIT-293 reagent (Mirus) for 293T and 293FT cells or Lipofectamine 2000 (Invitrogen) for U2OS cells following the manufacturer’s instructions. Lipofectamine RNAiMAX (Invitrogen) was used for siRNA transfections according to the manufacturer’s protocol.

##### Plasmids, shRNAs and antibodies

The ER_GFP__K20 construct were generated as previously reported. To generate lentiviral expression plasmids, the ER_GFP__K20 sequence were cloned into pLenti-CMV-GFP hygro (656-4) vector (Addgene #17446).

The human SAYSD1 cDNA was PCR-amplified from a human SAYSD1 cDNA clone (Origene) and inserted into the pLenti-cmv-GFP hygro (656-4) vector (Addgene #17446) to generate either N-terminally HA-tagged or C-terminally Myc-FLAG-tagged wildtype SAYSD1 (SAYSD1 WT) and N-terminal 17 residue-deleted SAYSD1 (*Δ*1-17 SAYSD1). The conserved “SAYSVFN” motif was mutated to “AAAAAAA” motif in the HA-SAYSD1 plasmid to generate the HA-SAYSD1 “7A” mutant. To generate a bacterial vector expressing human UFM1, the codon-optimized gene fragment encoding human UFM1 was inserted into the pET28a vector between NdeI and BamHI sites.

All shRNA lentiviral plasmids were generated based on the pLKO.1 puro vector (Addgene #8453) followed by the protocol in Addgene. The target sequences for shRNAs are listed below.

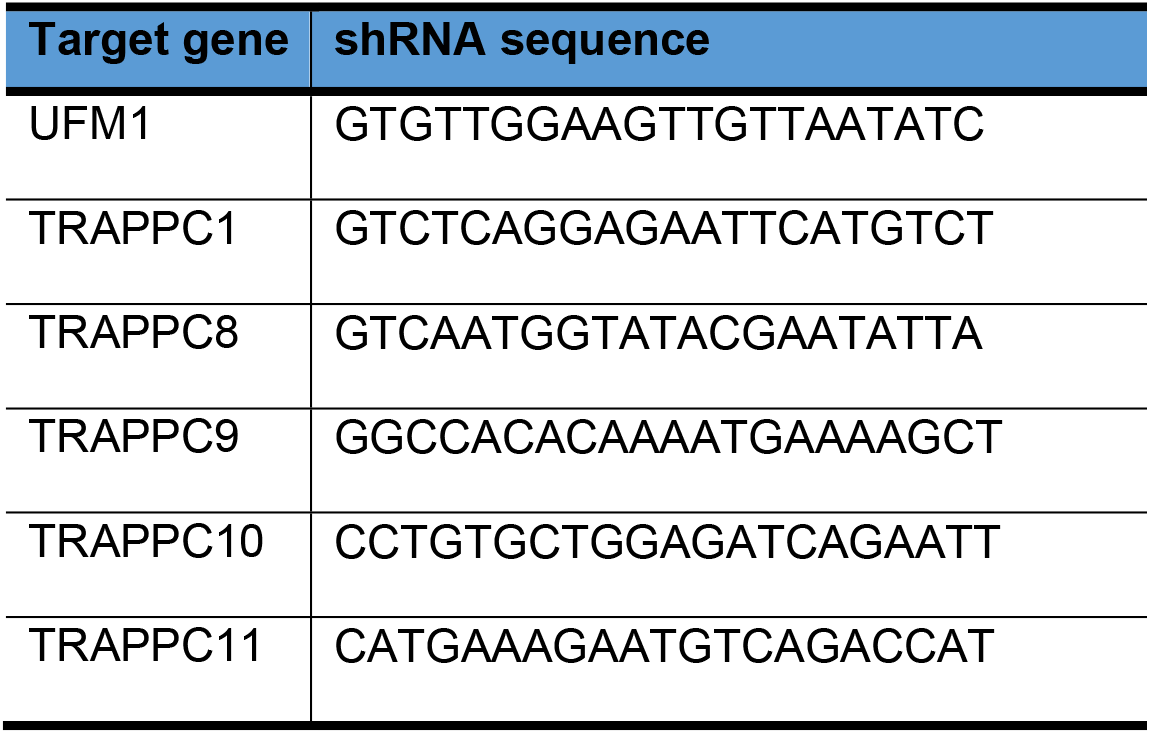

To generate GST-fusion proteins, the corresponding DNA fragments encoding different domains of SAYSD1 were either inserted into the pET41a vector to construct N-terminal GST fusion proteins or used to replace Mff(1-61) in the pET28-Mff(1-61)-PP-GST vector (Addgene #73042) to generate C-terminal GST-fusion proteins.

To generate transgenic flies expressing ER_GFP__K20, the pUAST-attb-ER_CFP__K20 plasmid was generated by inserting the ER_GFP__K20 sequence into the EcoRI and XbaI sites of pUAST-attb vector.

All plasmids were verified via Sanger sequencing and prepared using the Qiagen midiprep kit before use. Antibodies are listed in the resources table.

##### Lentivirus production and transduction

For the generation of human GeCKOv2 lentiviral pooled libraries, two 15-cm dishes of 293FT cells were seeded at 40% confluence. In the next day, 1 h prior to transfection, medium was replaced with 13 mL pre-warmed Opti-MEM medium (Life Technologies). For each dish, 6.8 μg pCMV-VSV-G, 10.1 μg psPAX2 (Addgene), 13 μg pLentiCRISPRv2-sgRNA plasmids pool and 135 μL PLUS reagent (Invitrogen) were added to 4mL Opti-MEM as mixture A, which is then mixed with mixture B containing 68 μL lipofectamine 2000 and 4mL Opti-MEM. The complete mixture was incubated for 20 min at RT before being added to cells. After 6 h, the medium was changed to 25 mL D10 medium (DMEM medium with 10% FBS and 1% Bovine Serum Albumin) with penicillin/streptomycin (10U/mL) for virus production. After 60 h of incubation, virus-containing medium from two culture dishes were combined and centrifuged at 3,000 rpm at 4 °C for 10 min and the supernatant was filtered through a 0.45 μm low protein-binding membrane (Steriflip HV/PVDF, Millipore). To concentrate lentivirus, the cleared supernatant was ultracentrifuged at 24,000 rpm for 2h at 4 °C using the JA25.50 rotor (Beckman). Virus was resuspended overnight in 180 μL D10 medium at 4 °C. Virus was aliquoted, flash-frozen in liquid nitrogen and stored at −80 °C.

For the shRNA lentivirus production, 293FT cells were seeded in a 6-well plate at 40% confluence. After 24h, cells in each well were transfected with 0.4 μg pCMV-VSV-G, 0.6 μg psPAX2 and 0.8 μg lentiviral shRNA plasmids using the TransIT-293 reagent (Mirus). After 6 h, the medium was replaced with fresh growth media and continued incubating for 60 h. To collect the lentivirus, the medium was filtered through a 0.45 μm PVDF membrane and used without further concentration.

##### Genome-wide CRISPR/Cas9 knockout screen

The GeckoV2 library was purchased from Addgene (1000000048) and amplified according to the Addgene’s protocol and the online protocol from Feng Zhang’s lab (Sanjana et al., 2014). The complexity of the sgRNA library was verified by high-throughput sequencing by the NIDDK Genomic Core. Lentiviruses containing the sgRNA library and Cas9 were generated and used for transduction via spinfection. Briefly, 60 million ER_GFP__K20 stable 293T cells supplemented with 8 μg/mL polybrene (Sigma) were seeded in two 12-well plates at a density of 3 million cells per well. Concentrated GeCKO v2 lentivirus was added to each well at a multiplicity of infection (MOI) at 0.3. Cells were then spun at 1000g at room temperature for 2h followed by incubation at 37 °C in a humidified incubator for 1h. After the medium was removed, fresh growth medium was added and incubated cells for 48 h before the start of selection for lentiviral integration using puromycin (0.3 μg/mL). The transduced cells were subcultured in medium supplemented with puromycin every 2 days for a total of 8 days and 80 million cells were maintained for each passage. After puromycin treatment, cells were recovered in a medium lacking puromycin for 24h before cell sorting. In total, 60 million cells were sorted into GFP high (1% of total cells) or GFP low (80% of total cells) cell populations by a FACS AriaII cell sorter. Genomic DNA was extracted from each cell population using a QIAGEN Blood Maxi kit (for GFP low cells) or QIAGEN Blood Midi kit (for GFP high cells) according to the manufacturer’s instructions. sgRNAs sequences were amplified from genomic DNA samples using the Herculase II Fusion DNA Polymerase (Agilent Technologies) in two PCR steps following the protocol as previously described. The resulting sgRNA libraries were sequenced using the Illumina NovaSeq at the NHLBI DNA Sequencing and Genomics core. sgRNA sequences were obtained per sample by extracting 20 bps followed by the index sequence of “TTGTGGAAAGGACGAAACACCG” on the de-multiplexed FASTQ files from Illumina’s NGS sequencer using the Cutadapt software, version 2.8 (https://doi.org/10.14806/ej.17.1.200). FASTQC, version 0.11.9 was used to assess the sequencing quality (http://www.bioinformatics.babraham.ac.uk/projects/fastqc/). MAGeCK, version 0.5.9, was used to quantify and to identify differentially expressed sgRNAs (Li et al., 2014). MAGeCK count command was run on the merged two-half libraries of A and B to quantify. Differentially expressed sgRNAs with statistical significance were determined by running the MAGeCK test command in paired mode. Genes were ranked based on the number of unique sgRNA enriched in the GFP high population versus the GFP low population. Data was derived from two biological repeats of the screen.

##### Immunofluorescence microscopy and live-cell imaging

293T or COS7 cells were co-transfected with LAMP1-mCherry and either ER_GFP__K20 or YFP-PrP* (C179A) using Lipofectamine 3000. Medium was changed after 16h and cells were seeded onto a glass coverslip. For 293T cells only, coverslips were pre-coated for 3 h with 0.01% poly-L-lysine solution to promote adherence to the coverslip. Forty-eight hours after transfection, cells were incubated with 0.4 mg/mL Alexa Fluor 647-conjugated anti-GFP antibody and 250 nM Bafilomycin A1, and imaged every 30 min for 15 h or more, as indicated. To detect antibody uptake of Alexa Fluor 647-conjugated anti-GFP antibody, image acquisition in the 647 nm channel from ER_GFP__K20-expressing or YFP-PrPC179-expressing cells were identical in every way, including using identical excitation, emission, gain and contrast parameters. The imaging parameters were set to detect ER_GFP__K20 or YFP-PrP* at steady-state (t=0). While imaging ER_GFP__K20 or YFP-PrP* in Bafilomycin A1 treated cells, ER_GFP__K20 or YFP-PrP* quickly saturated the pixels due to the rapid rate of accumulation in lysosomes and limitations in the dynamic range of our camera.

Images and time lapse series were collected with Nikon inverted spinning disk confocal microscope equipped with Yokogawa CSU-X1 Spinning Disk, EMCCD camera, laser launch including 488 nm (to image GFP), 561 nm (to image mCherry) and 647 nm (to image Alexa Fluor 647), focus drift correction through the Perfect Focus System, 60x Plan Apo 1.40 NA oil / 0.13mm WD, and controlled by the NIS-Elements software. For live cell imaging experiments, cells seeded in # 1.5 coverslip bottom-dishes and incubated in complete culture medium (described above) at 37°C with 5% CO2 within a Tokai incubator on the microscope stage. Imaging experiments were performed at 60-80% confluency 48 h after transient transfection.

For fixed cells imaging, cells were fixed in 4% paraformaldehyde at room temperature for 12 min followed by washing with PBS for two times. The cells were permeabilized in PBS containing 0.1% Triton X-100 and 10% FBS at room temperature for 45 min. Fixed cells were then incubated with respective primary antibodies at indicated concentration at 4 °C overnight, followed by incubation with a secondary antibody at room temperature for 1h. DAPI was used to stain nuclei. The cells were then imaged either using the Zeiss LSM 780 confocal microscope or the Nikon CSU-W1 SoRa microscope.

##### Immunoblotting

To perform protein immunoblotting, proteins were separated in NuPAGE (4%–12%) Bis-Tris gels (Invitrogen) and transferred onto nitrocellulose membranes (BioRad). Target protein was detected by specific primary antibodies followed by secondary horseradish peroxidase (HRP)-conjugated antibodies or fluorescently labeled secondary antibodies (Invitrogen). Enhanced chemiluminescence (ECL) method was used to develop protein signals immunoblotted with HRP-conjugated secondary antibodies. Immunoblotting signal was detected using the BioRad ChemiDoc Imaging System. The intensity of the detected protein bands was quantified by the BioRad Image Lab software.

##### Generation of SAYSD1 CRISPR knockout cells

To generate SAYSD1 knockout cells, a pLentiCRISPRv2 plasmid containing a SAYSD1-targeting sgRNA sequence (CTCAGCTAACCGCTGTTCCA) was generated and packed into lentivirus. 293T cells transduced by the lentivirus were selected with puromycin for 7 days and the surviving cells were single-cell sorted into 96-well plate. Clones were screened by immunoblotting using the anti-SAYSD1 antibody. Knockout clones were pooled and used for the experiments.

##### Procollagen I secretion

U2OS cells were seeded at 1.5 million cells per well in a 6-well plate supplemented with 50 μg/mL L-ascorbate acid. Twenty hours later, cells were washed once and incubated in 1.2 mL culture media supplemented with 50 μg/mL L-ascorbate acid and 50 μg/mL cycloheximide (CHX). The medium was collected at 0, 1, 2 and 3 h time points, centrifuged at 2000 g for 5 min to remove any cells or cell debris. To prepare cell lysate, cells were washed once with ice-cold PBS and lysed in 90 μL NP40 lysis buffer and incubated for 15 min on ice, followed by centrifugation at 15.6 kg for 10 min. The medium and lysate were diluted 10 or 50 times, respectively, and the procollagen I level in medium and cell lysate was measured by the human Procollagen I alpha 1 ELISA Kit from Abcam.

To assess the effect of Baf A1 treatment on procollagen I secretion, U2OS cells were treated with DMSO or 250 nM Baf A1 for 5 hours and the procollagen I level in medium and cell lysate were measured as above-mentioned.

##### Quantitative Reverse transcription PCR (qRT-PCR)

Total cellular RNA was extracted using the TriPure reagent (Roche) and purified using the RNeasy MinElute Cleanup Kit (Promega) following standard protocols. RNA concentration was measured by a Nanodrop UV spectrophotometer, and 500 ng RNA was used to prepare cDNA library using the BioRad iScript Reverse Transcription Supermix. The qPCR reactions were carried out using the BioRad SsoAdvanced SYBR Green Supermix kit on a BioRad CFX96 machine. Data was analyzed using BioRad CFX manager 3.0 software. Ribosomal 18S RNA was used as a reference gene for quantification of gene expression. The following primers were used for qPCR: *COPB1:* forward, 5’-TTAGCTTAAAAAATGATCTAG-3’; reverse, 5’-GAAAACGAAGAGTAGATCCTC-3’; *KDELR2:* forward, 5’-CTGCTGAAGATCTGGAAGAC-3’; reverse, 5’-AGAGGATCTCAAGAGGAGAG-3’; *RAB1B:* forward, 5’-CCTGCTCCTGCGGTTTGCTG-3’; reverse, 5’-TCTTGTTGCCCACCAGGAGC-3’; *TMED3:* forward, 5’-AGTTCTCCCTGGATTACCAG-3’; reverse, 5’-CCTCATGGATGGTCACGCAG-3’; *TRAPPC1:* forward, 5’-GAAGCAAGCAGGGATTCCCAA-3’; reverse, 5’-TAGATGTGGTGCAGCACATCT-3’; *TRAPPC8*: forward, 5’-AGCAGGAGATTATGACCTTAA-3’; reverse, 5’-GTATTTAAGTGTATTTGGTAT-3’; *TRAPPC9*: forward, 5’-GAGCTTCACAGAGGAAGTGAA-3’; reverse, 5’-AGTGGTGCCTGTAGCGGATGT-3’; *TRAPPC10*: forward, 5’-CTTCCAAGAGAACCAATGGAA-3’; reverse, 5’-CTTCAGAACATTCTGCCACTT-3’; *TRAPPC11:* forward, 5’-CCAAATGTAGACCCAAGAGAA-3’; reverse, 5’-GAACCACTGCAACTTTTGTGT-3’; *COL1A1*, forward, 5’-GATTCCCTGGACCTAAAGGTGC-3’; reverse, 5’-AGCCTCTCCATCTTTGCCAGCA-3’.

##### Radiolabeling pulse-chase assay

Radiolabeling pulse-chase assay was performed as previously described (Wang et al., 2020). To measure the turnover of ER_GFP__K20, 4.0 million 293T cells stably expressing ER_GFP__K20 or in some cases cells transfected with ER_GFP__K20 were starved in 2 mL starvation medium (DMEM medium lacking cysteine and methionine, supplemented with 10% FBS) for 30 min at 37 °C, followed by labeling with 4 mCi [35S]-methionine/cysteine in 300 μL starvation medium for 16 min at 37 °C. After labeling, cells were resuspended in 300 μL starvation medium supplemented with 2.5 mM methionine and cysteine. An aliquot of cells (65 μL) was immediately taken out and mixed with 500 μL RIPA buffer with 1 mM DTT and protease inhibitors. The remaining cells were incubated at 37 °C for 20, 40, and 60 minutes. Equal amounts of cells (65 μL) were taken at each time points and lysed in RIPA buffer. After centrifugation of cell lysate at 14,000× g for 5 min, the cleared cell extracts were subjected to immunoprecipitation using anti-GFP antibody and protein A beads. The beads were then washed twice with the NET buffer with 0.1% SDS and bound proteins were eluted in 36 μL Laemmli sample buffer by boiling at 95 °C for 5 min. Eluted proteins were analyzed by the SDS-PAGE using Criterion™ TGX™ precast gels (10%) followed by autoradiography analyses.

##### Ribosome fractionation

293T cells were seed into two wells at 0.5 million per well. Two wells were used for each siRNA transfection. siRNA transfections were performed twice on the following consecutive two days, with 60 pmol scrambled or UFM1 siRNA for each transfection. At 48 hours post-transfection, cells from two wells were combined and seeded into a 10-cm dish. One day later, cells were then transfected with 5 μg plasmid encoding ER_GFP__K20. The cells were then harvested 24 h later and used for radiolabeling pulse-chase as described above. Cells were chased for 0, 5, 10 and 20 min and 2 million cells were lysed at each time point in 200 μL polysome lysis buffer (25 mM Tris-HCl, pH 7.4, 150mM NaCl, 15mM MgCl_2_, 1 mM DTT, 1% Triton X-100, 200U/mL SUPERase In RNase inhibitor, 20 U/mL Turbo DNase and complete EDTA-free protease inhibitor cocktail) and incubated on ice for 10 min followed by centrifugation at 14,000 g for 5 min. Two hundred microliter supernatant was loaded onto a sucrose cushion (0.8 M sucrose, 1 mM DTT, 200 U/mL SUPERase In RNase inhibitor and complete EDTA-free protease inhibitor cocktail) and centrifuged at 69000 rpm using a TLA100.4 rotor for 2.5 hours at 4 °C. After the ribosome-free supernatant was removed, the ribosome pellet was washed once with water and dissolved in 500 μL RIPA buffer. After centrifugation at 14000 g for 5 min, the cleared ribosome solutions and the ribosome-free fractions were used for immunoprecipitation using anti-FLAG beads. Samples were resolved using SDS-PAGE and analyzed by autoradiograph.

To examine the association of SAYSD1 with Sec61β and ribosomes, 293T cells plated in a 10 cm dish were transfected with 5 μg ER_GFP__K20 plasmid. 72 hours post transfection, cells were lysed by 1% CHAPS in buffer LC (20 mM Tris-HCl, pH 8.0, 150 mM NaCl, 15 mM MgCl2, 1 mM DTT) that also contains 100 μg/mL cycloheximide, 200 U/mL SUPERase In RNase inhibitor, 20 U/mL Turbo DNase and a complete EDTA-free protease inhibitor cocktail. Cleared cell lysate (12,000 g 10min) was layered on top of a 10-50% sucrose gradient in the buffer LC containing 0.1 % CHAPS. The sample was centrifuged at 44,000 rpm for 3 h in a SW60 rotor. 500 μL fractions were collected for immunoblotting analysis.

##### Immunoprecipitation

To detect interaction of endogenous SAYSD1 with the translocon and ribosomes. Four million of *SAYSD1::GFP* 293T cells or parental cells were seed in 10-cm dishes. One day later, cells were treated with 200 nM anisomycin (ANS) or DMSO for 1 or 2 hours and lysed in 600 ul of digitonin lysis buffer (25 mM HEPES, pH 7.3, 250 mM KOAc, 10 mM MgCl_2_, 5mM NaOAc and 0.5 mM EGTA, 2% digitonin, 1 mM DTT and complete protease inhibitors) at 4°C for 15 min. Sucrose was added to 250 mM and cell lysate was centrifuged at 15.6 kg for 5 min. For immunoprecipitation, 0.5 mL cleared cell lysate was incubated with 30 uL slurry of pre-washed GFP-Trap nanobeads (ChromoTek) for 1.5 h at 4°C. After incubation, beads were washed with digitonin wash buffer (25 mM HEPES, pH 7.3, 115 mM KOAc, 10 mM Mg KOAc, 5mM NaOAc, 0.2% digitonin, and 0.5 mM EGTA) followed by boiling in 50 μL Laemmli sample buffer at 95 °C for 10 min before SDS-PAGE and immunoblotting analyses. To immunoprecipitate the translocon, 293T cells were lysed in the digitonin lysis buffer and cleared supernatant was incubated with 10 μL pre-immune or Sec61β serum for 1h at 4°C. Protein A beads were used to precipitate antibody protein complexes

To detect interaction of TAQC substrates with SAYSD1, 293T cells were seeded in a 6-well plate at 0.5 million per well and grown for 24 h followed by transfection of 1.5 μg ER_GFP__K20-expressing plasmid. For control experiments, cells were transfected with ER_K0 or pcDNA3 empty vectors. After 48 hours, cells were washed with ice-cold PBS and lysed in 0.5 mL CHAPS lysis buffer (1% CHAPS, 50 mM HEPES pH7.3, 100 mM NaCl, 1 mM DTT and complete protease inhibitors) for 15 min at 4°C. The cell extracts were then centrifuged at 15.6 kg for 10 min at 4 °C. To immunoprecipitate ER_GFP__k20, the cleared supernatant was incubated with 30 μL preequilibrated anti-FLAG M2 beads at 4°C for 1 h using a head-over-tail rotator. For GST-N17 pulldown of ribosome, 2 x 10^6^ cells were lysed in 0.4 mL CHAPS lysis buffer. The cleared supernatant was adjusted to include 1mM EGTA and 2.5mM MgCl_2_ and then incubated with glutathione beads prebound with GST-tagged recombinant proteins for 30 min at 4°C. After incubation, beads were washed twice with CHAPS wash buffer (0.02% CHAPS, 50 mM HEPES pH7.3, 100 mM NaCl, 1 mM DTT) and bound proteins were eluted by boiling in 50 μL Laemmli sample buffer at 95 °C for 10 min. Samples were analyzed by SDS-PAGE and immunoblotting using respective antibodies as indicated.

To detect the interaction of GST-SAYSD1 with UFM1, GST-tagged proteins (2 μg) and His_6_-UFM1 or Atto565-UFM1 (16 μg) were mixed in 200 μL UFM1 binding buffer (25 mM of Tris-HCl, pH 7.4, 150 mM of NaCl, 2 mM of MgCl_2_, 2 mM of KCl, and 0.05% of NP40). After incubation at 4 °C for one hour, the mixture was transferred into a new microtube containing 25 μL (bed volume) GST beads equilibrated with the UFM1 binding buffer and further incubated for another hour. The beads were spun down at 1,000 g 1min and washed twice with the UFM1 binding buffer. The proteins on beads were eluted with 50 μL of Laemmli sample buffer and assayed by SDS-PAGE followed by immunoblotting or fluorescence scanning.

##### Endogenous tagging

GFP tagging of endogenous SAYSD1 was carried out using a CRISPR/Cas12-Assisted PCR Tagging system as described (http://www.pcr-tagging.com) (Fueller et al., 2020). Briefly, the PCR cassette was amplified from pMaCTag-05 plasmid by the AccuPrime™ Pfx DNA Polymerase with the primers, M1_SAYSD1 and M2_SAYSD1, listed below. The PCR product was gel purified with QIAGEN Gel Extraction Kit. 293T cells were transiently transfected with 1 μg of the PCR cassette and 1 μg of pcDNA3.1-hAsCpf1/TYCV/pY210 (gift from Feng Zhang, Addgene plasmid # 89351) using TransIT293 (Mirus) according to the manufacturing protocol and GFP positive cells were sorted two weeks later with Flow Cytometry.

M1_SAYSD1: 5’-GTGTTCAATCCAGGCTGTGAAGCCATCCAGGGCACCCTGACTGCAGAGCAGTTGGAGCGCGAGTTACAGTTGAGACCCCTGGCAGGGAGATCAGGTGGAGGAGGTAGTG-3’M2_SAYSD1: 5’-CCAATGGTGAGGAAGACCACATCAGAGGTTAGCTGCATGACAGCACAGCTGGGTCAAAAAAGGGTCCTATCTCCCTGCCAGATCTACAAGAGTAGAAATTAGCTAGCTGCATCGGTACC-3’

##### Recombinant protein purification and mass photometry

Plasmids encoding the N-terminally His_6_-tagged UFM1 were transformed into BL21(DE3) cells (Agilent) and the transformed cells were cultured in LB medium supplemented with kanamycin 50 μg/mL and chloramphenicol 25 μg/mL. To induce protein expression, 0.5 mM IPTG were added when the optical density at 600 nm of the bacterial culture reached 0.9, and the proteins are expressed for 22 h at 19 °C. For protein purification, bacterial pellet was lysed in buffer R (20 mM Tris pH 8.0, 500 mM KCl, 10% glycerol) supplemented with the EDTA-free complete protease inhibitor cocktail by sonication and centrifuged for 45 min at 15000 rpm using a JA15.25 rotor. The cleared supernatant was incubated with Ni-NTA agarose (Qiagen) for 1 h at 4 °C. After washing, the His_6_-UFM1 protein was eluted in buffer R supplemented with 50-300 mM imidazole. Elution fractions were analyzed using SDS-PAGE followed by Coomassie blue staining. The proteins are pooled, concentrated and dialyzed in a protein buffer (50 mM Tris, pH 7.4, 50 mM NaCl). GST-fusion proteins were expressed following the abovementioned procedure and purified using the Glutathione Sepharose 4B resin using the standard protocol. Proteins were then dialyzed against assay buffer. Protein concentration was determined by the Bradford assay.

The isolation of ribosomes from rabbit reticulocyte lysate (RRL) was performed according to a published protocol (Rivera et al., 2015). Briefly, in a fresh ultracentrifuge tube, 250 μL RRL was laid on 1 mL of a cushion buffer (50 mM Tris-HCl, pH7.5, 5mM MgCl2, 25 mM KCl, 2 M Sucrose, filter-sterilized through a 0.22 mm filter). The sample was placed in a TLA120.2 rotor, and centrifuged at 4 °C at 48,000 rpm, for 20 hours in Beckman OptimaTM MAX ultracentrifuger. After centrifuge, the supernatant was carefully removed, and the ribosome-containing pellet was resuspended in 250 μL PBS buffer.

Mass photometry was performed on an OneMP instrument (Refeyn, UK) at room temperature following the published protocol (Wu and Piszczek, 2020). Briefly, microscope coverslips (24 × 50 mm, Fisher Scientific) were rinsed consecutively in isopropanol and H2O and blow dry in a stream of clean nitrogen. 30nM GST or MH-GST were mixed with the indicated concentrations of UFM1 in a phosphate saline buffer. After applying 10 μl solution on the slide, the mass distribution plot was obtained with the software provided by the instrument manufacturer (Refeyn, UK)

##### Fly experiments

Tissue dissection and immunostaining were performed as described previously (Ye and Fortini, 1998). To examine COL4 biogenesis, Viking-GFP was recombined to the same chromosome as Cg-Gal4 in female germlines. Progenies were screened based on eye color. Cg-Gal4, Viking-GFP females were crossed to w^1118^/y; UAS-UFM1 shRNA/UAS-mCherry or UAS-Control shRNA/UAS-mCherry. Third instar larvae without UAS-mCherry were picked for dissection, fixing, and confocal imaging. For phalloidin staining, tissues were fixed in 4% formaldehyde in phosphate buffer saline (PBS) at room temperature for 15 min, permeabilized in PBS with 0.1% Saponin and 10% fetal bovine serum and stained by Acti-stain™ 555 Phalloidin at 50 nM for 30 min at room temperature.

To generate SAYSD1 CRISPR KO flies, female flies expressing sgRNA ubiquitously for Cas9-mediated mutagenesis of CG13663 (FBgn0039291) (#82685) were crossed to flies expressing Cas9 in the male germline cells with the Nos promoter (Port et al., 2014). Male F1 progenies bearing both the sgRNA-expressing chromosome and the Nos-Cas9 chromosome were then crossed to female flies bearing double balancers for the second and third chromosome (x/y, w^-^; *Sp/Cyo; TM2/TM6B*). Individual F2 male progenies with white eyes were then crossed to flies with the double balancers to establish candidate KO lines. Genomic DNA from homozygous SAYSD1 mutant was extracted and used to amplify the SAYSD1 genomic DNA and for sequencing validation.

##### Quantification and statistical analyses

Immunoblotting plots were digitally scanned by a ChemDoc Imaging System. Band intensity was measured by Image Lab (BioRad). Imaging data were processed by either Zen 2.3 SP1 (Zeiss) or by Nikon Elements (Nikon). Fluorescence intensity was measured by Imaged J or by Imaris in the case of 3D imaging. Unless specified, the n values in the figure legend indicate the number of independent biological replicates. Statistically analyses were performed using GraphPad Prism 9. For two group comparison, unpaired Student’s t-test was used. For multi-group comparison, we used one-way ANOVA. For comparing life span in flies, Gehan-Breslow-Wilcoxon test was used. Error bars, shown as mean±SD and p-values were calculated by GraphPad Prism 9. A value of p < 0.05 was considered as statistically significant. Graphs were prepared with GraphPad Prism 7.

##### A list of significant gene hits from the CRISPR screen can be found in Table S1

**Supplemental movie 1** A time lapse video shows HEK293T cells transfected with ER_GFP__K20 and treated with Baf A1 and a Lysotracker red dye.

## Supporting information

Supplemental Figure 1-7 with legends

## Acknowledgments

We thank J. Skeath (Washington University) for Viking-GFP flies, H. Ryoo (New York University) for the Xbp1-GFP fly strain, the NHLBI Genomic, flow cytometry, Biophysics, and the NIDDK Advanced Light Microscopy & Image Analysis Cores (ALMIAC) for services, N. Guydosh (NIDDK) and J. Yewdell (NIAID) for critical reading of the manuscript. This research was supported by an intramural research program of the NIDDK (Y. Ye) and of NINDS (Y. Quan) in the National Institutes of Health, and by NIH/NIGMS R01 GM134327 to P. Satpute-Krishnan.

## Author contributions

L. W. designed and performed the CRISPR/Cas9 screen and identified SAYSD1. X. Y. made stable cell lines, performed endogenous tagging, imaging experiments, and *in vitro* pulldown experiments. S. Yun (NIDDK) analyzed the CRISPR screen data, Q. Y. provided fly strains and advised on the fly study. P. S.-K. analyzed the trafficking path for ER_GFP__K20. Y.Y. performed the fly experiments and oversaw the study. L. *W*. and Y.Y. wrote the paper.

## References

Ast, T., Michaelis, S., and Schuldiner, M. (2016). The Protease Ste24 Clears Clogged Translocons. Cell 164, 103–114.

Barrowman, J., Bhandari, D., Reinisch, K., and Ferro-Novick, S. (2010). TRAPP complexes in membrane traffic: convergence through a common Rab. Nat Rev Mol Cell Biol 11, 759–763.

Bengtson, M.H., and Joazeiro, C.A. (2010). Role of a ribosome-associated E3 ubiquitin ligase in protein quality control. Nature 467, 470–473.

Bienkowski, R.S. (1989). Intracellular degradation of newly synthesized collagen. Revis Biol Celular 21, 423–443.

Cai, Y., Pi, W., Sivaprakasam, S., Zhu, X., Zhang, M., Chen, J., Makala, L., Lu, C., Wu, J., Teng, Y., et al. (2015). UFBP1, a Key Component of the Ufm1 Conjugation System, Is Essential for Ufmylation-Mediated Regulation of Erythroid Development. PLoS Genet 11, e1005643.

Colin, E., Daniel, J., Ziegler, A., Wakim, J., Scrivo, A., Haack, T.B., Khiati, S., Denomme, A.S., Amati-Bonneau, P., Charif, M., et al. (2016). Biallelic Variants in UBA5 Reveal that Disruption of the UFM1 Cascade Can Result in Early-Onset Encephalopathy. Am J Hum Genet 99, 695–703.

Duan, R., Shi, Y., Yu, L., Zhang, G., Li, J., Lin, Y., Guo, J., Wang, J., Shen, L., Jiang, H., et al. (2016). UBA5 Mutations Cause a New Form of Autosomal Recessive Cerebellar Ataxia. PLoS One 11, e0149039.

Forrester, A., De Leonibus, C., Grumati, P., Fasana, E., Piemontese, M., Staiano, L., Fregno, I., Raimondi, A., Marazza, A., Bruno, G., et al. (2019). A selective ER-phagy exerts procollagen quality control via a Calnexin-FAM134B complex. EMBO J 38.

Fueller, J., Herbst, K., Meurer, M., Gubicza, K., Kurtulmus, B., Knopf, J.D., Kirrmaier, D., Buchmuller, B.C., Pereira, G., Lemberg, M.K., et al. (2020). CRISPR-Cas12a-assisted PCR tagging of mammalian genes. J Cell Biol 219.

Gelse, K., Poschl, E., and Aigner, T. (2003). Collagens--structure, function, and biosynthesis. Adv Drug Deliv Rev 55, 1531–1546.

Gerakis, Y., Quintero, M., Li, H., and Hetz, C. (2019). The UFMylation System in Proteostasis and Beyond. Trends Cell Biol 29, 974–986.

Gregorio, I., Braghetta, P., Bonaldo, P., and Cescon, M. (2018). Collagen VI in healthy and diseased nervous system. Dis Model Mech 11.

Huter, P., Arenz, S., Bock, L.V., Graf, M., Frister, J.O., Heuer, A., Peil, L., Starosta, A.L., Wohlgemuth, I., Peske, F., et al. (2017). Structural Basis for Polyproline-Mediated Ribosome Stalling and Rescue by the Translation Elongation Factor EF-P. Mol Cell 68, 515–527 e516.

Ito, S., and Nagata, K. (2021). Quality Control of Procollagen in Cells. Annu Rev Biochem 90, 631–658.

Ito-Harashima, S., Kuroha, K., Tatematsu, T., and Inada, T. (2007). Translation of the poly(A) tail plays crucial roles in nonstop mRNA surveillance via translation repression and protein destabilization by proteasome in yeast. Genes Dev 21, 519–524.

Joazeiro, C.A.P. (2017). Ribosomal Stalling During Translation: Providing Substrates for Ribosome-Associated Protein Quality Control. Annu Rev Cell Dev Biol 33, 343–368.

Jumper, J., Evans, R., Pritzel, A., Green, T., Figurnov, M., Ronneberger, O., Tunyasuvunakool, K., Bates, R., Zidek, A., Potapenko, A., et al. (2021). Highly accurate protein structure prediction with AlphaFold. Nature.

Juszkiewicz, S., and Hegde, R.S. (2017). Initiation of Quality Control during Poly(A) Translation Requires Site-Specific Ribosome Ubiquitination. Mol Cell 65, 743–750 e744.

Kang, S.H., Kim, G.R., Seong, M., Baek, S.H., Seol, J.H., Bang, O.S., Ovaa, H., Tatsumi, K., Komatsu, M., Tanaka, K., et al. (2007). Two novel ubiquitin-fold modifier 1 (Ufm1)-specific proteases, UfSP1 and UfSP2. J Biol Chem 282, 5256–5262.

Kramer, G., Rauch, T., Rist, W., Vorderwulbecke, S., Patzelt, H., Schulze-Specking, A., Ban, N., Deuerling, E., and Bukau, B. (2002). L23 protein functions as a chaperone docking site on the ribosome. Nature 419, 171–174.

Lemaire, K., Moura, R.F., Granvik, M., Igoillo-Esteve, M., Hohmeier, H.E., Hendrickx, N., Newgard, C.B., Waelkens, E., Cnop, M., and Schuit, F. (2011). Ubiquitin fold modifier 1 (UFM1) and its target UFBP1 protect pancreatic beta cells from ER stress-induced apoptosis. PLoS One 6, e18517.

Li, W., Xu, H., Xiao, T., Cong, L., Love, M.I., Zhang, F., Irizarry, R.A., Liu, J.S., Brown, M., and Liu, X.S. (2014). MAGeCK enables robust identification of essential genes from genome-scale CRISPR/Cas9 knockout screens. Genome Biol 15, 554.

Liang, J.R., Lingeman, E., Luong, T., Ahmed, S., Muhar, M., Nguyen, T., Olzmann, J.A., and Corn, J.E. (2020). A Genome-wide ER-phagy Screen Highlights Key Roles of Mitochondrial Metabolism and ER-Resident UFMylation. Cell 180, 1160–1177 e1120.

Morin, X., Daneman, R., Zavortink, M., and Chia, W. (2001). A protein trap strategy to detect GFP-tagged proteins expressed from their endogenous loci in Drosophila. Proc Natl Acad Sci U S A 98, 15050–15055.

Muona, M., Ishimura, R., Laari, A., Ichimura, Y., Linnankivi, T., Keski-Filppula, R., Herva, R., Rantala, H., Paetau, A., Poyhonen, M., et al. (2016). Biallelic Variants in UBA5 Link Dysfunctional UFM1 Ubiquitin-like Modifier Pathway to Severe Infantile-Onset Encephalopathy. Am J Hum Genet 99, 683–694.

Nahorski, M.S., Maddirevula, S., Ishimura, R., Alsahli, S., Brady, A.F., Begemann, A., Mizushima, T., Guzman-Vega, F.J., Obata, M., Ichimura, Y., et al. (2018). Biallelic UFM1 and UFC1 mutations expand the essential role of ufmylation in brain development. Brain 141, 1934–1945.

Palacios-Macapagal, D., Cann, J., Cresswell, G., Zerrouki, K., Dacosta, K., Wang, J., Connor, J., and Davidso, T.S. (2021). Chronic CNS Pathology is Associated with Abnormal Collagen Deposition and Fibrotic-like Changes. Biorxiv.

Pastor-Pareja, J.C., and Xu, T. (2011). Shaping cells and organs in Drosophila by opposing roles of fat body-secreted Collagen IV and perlecan. Dev Cell 21, 245–256.

Peil, L., Starosta, A.L., Lassak, J., Atkinson, G.C., Virumae, K., Spitzer, M., Tenson, T., Jung, K., Remme, J., and Wilson, D.N. (2013). Distinct XPPX sequence motifs induce ribosome stalling, which is rescued by the translation elongation factor EF-P. Proc Natl Acad Sci U S A 110, 15265–15270.

Phillips, B.P., and Miller, E.A. (2020). Ribosome-associated quality control of membrane proteins at the endoplasmic reticulum. J Cell Sci 133.

Port, F., Chen, H.M., Lee, T., and Bullock, S.L. (2014). Optimized CRISPR/Cas tools for efficient germline and somatic genome engineering in Drosophila. Proc Natl Acad Sci U S A 111, E2967–2976.

Qin, B., Yu, J., Nowsheen, S., Wang, M., Tu, X., Liu, T., Li, H., Wang, L., and Lou, Z. (2019). UFL1 promotes histone H4 ufmylation and ATM activation. Nat Commun 10, 1242.

Rapoport, T.A., Li, L., and Park, E. (2017). Structural and Mechanistic Insights into Protein Translocation. Annu Rev Cell Dev Biol 33, 369–390.

Reggio, A., Buonomo, V., Berkane, R., Bhaskara, R.M., Tellechea, M., Peluso, I., Polishchuk, E., Di Lorenzo, G., Cirillo, C., Esposito, M._✓_ et al. (2021). Role of FAM134 paralogues in endoplasmic reticulum remodeling, ER-phagy, and Collagen quality control. EMBO Rep 22, e52289.

Rivera, M.C., Maguire, B., and Lake, J.A. (2015). Purification of polysomes. Cold Spring Harb Protoc 2015, 303–305.

Ryan, M.D., King, A.M., and Thomas, G.P. (1991). Cleavage of foot-and-mouth disease virus polyprotein is mediated by residues located within a 19 amino acid sequence. J Gen Virol 72 (Pt 11), 2727–2732.

Sacher, M., Jiang, Y., Barrowman, J., Scarpa, A., Burston, J., Zhang, L., Schieltz, D., Yates, J.R., 3rd, Abeliovich, H., and Ferro-Novick, S. (1998). TRAPP, a highly conserved novel complex on the cis-Golgi that mediates vesicle docking and fusion. EMBO J 17, 2494–2503.

Sanjana, N.E., Shalem, O., and Zhang, F. (2014). Improved vectors and genome-wide libraries for CRISPR screening. Nat Methods 11, 783–784.

Sapperstein, S.K., Walter, D.M., Grosvenor, A.R., Heuser, J.E., and Waters, M.G. (1995). p115 is a general vesicular transport factor related to the yeast endoplasmic reticulum to Golgi transport factor Uso1p. Proc Natl Acad Sci U S A 92, 522–526.

Satpute-Krishnan, P., Ajinkya, M., Bhat, S., Itakura, E., Hegde, R.S., and Lippincott-Schwartz, J. (2014). ER stress-induced clearance of misfolded GPI-anchored proteins via the secretory pathway. Cell 158, 522–533.

Shao, S., Brown, A., Santhanam, B., and Hegde, R.S. (2015). Structure and assembly pathway of the ribosome quality control complex. Mol Cell 57, 433–444.

Stephani, M., Picchianti, L., and Dagdas, Y. (2021). C53 is a cross-kingdom conserved reticulophagy receptor that bridges the gap betweenselective autophagy and ribosome stalling at the endoplasmic reticulum. Autophagy 17, 586–587.

Sun, Z., and Brodsky, J.L. (2019). Protein quality control in the secretory pathway. J Cell Biol 218, 3171–3187.

Tatsumi, K., Yamamoto-Mukai, H., Shimizu, R., Waguri, S., Sou, Y.S., Sakamoto, A., Taya, C., Shitara, H., Hara, T., Chung, C.H., et al. (2011). The Ufm1-activating enzyme Uba5 is indispensable for erythroid differentiation in mice. Nat Commun 2, 181.

Voorhees, R.M., Fernandez, I.S., Scheres, S.H., and Hegde, R.S. (2014). Structure of the mammalian ribosome-Sec61 complex to 3.4 A resolution. Cell 157, 1632–1643.

Walczak, C.P., Leto, D.E., Zhang, L., Riepe, C., Muller, R.Y., DaRosa, P.A., Ingolia, N.T., Elias, J.E., and Kopito, R.R. (2019). Ribosomal protein RPL26 is the principal target of UFMylation. Proc Natl Acad Sci U S A 116, 1299–1308.

Wang, L., Xu, Y., Rogers, H., Saidi, L., Noguchi, C.T., Li, H., Yewdell, J.W., Guydosh, N.R., and Ye, Y. (2020). UFMylation of RPL26 links translocation-associated quality control to endoplasmic reticulum protein homeostasis. Cell Res 30, 5–20.

Wang, L., and Ye, Y. (2020). Clearing Traffic Jams During Protein Translocation Across Membranes. Front Cell Dev Biol 8, 610689.

Wegrzyn, R.D., Hofmann, D., Merz, F., Nikolay, R., Rauch, T., Graf, C., and Deuerling, E. (2006). A conserved motif is prerequisite for the interaction of NAC with ribosomal protein L23 and nascent chains. J Biol Chem 281, 2847–2857.

Wu, D., and Piszczek, G. (2020). Measuring the affinity of protein-protein interactions on a single-molecule level by mass photometry. Anal Biochem 592, 113575.

Ye, Y., and Fortini, M.E. (1998). Characterization of Drosophila Presenilin and its colocalization with Notch during development. Mech Dev 79, 199–211.

Yurchenco, P.D., and Furthmayr, H. (1984). Self-assembly of basement membrane collagen. Biochemistry 23, 1839–1850.

Zhang, Y., Zhang, M., Wu, J., Lei, G., and Li, H. (2012). Transcriptional regulation of the Ufm1 conjugation system in response to disturbance of the endoplasmic reticulum homeostasis and inhibition of vesicle trafficking. PLoS One 7, e48587.

