## Supplemental Figure 1-7 with legends for "SAYSD1 senses UFMylated ribosome to safeguard co-translational protein translocation at the endoplasmic reticulum"

**Supplementary Figures and Legends**

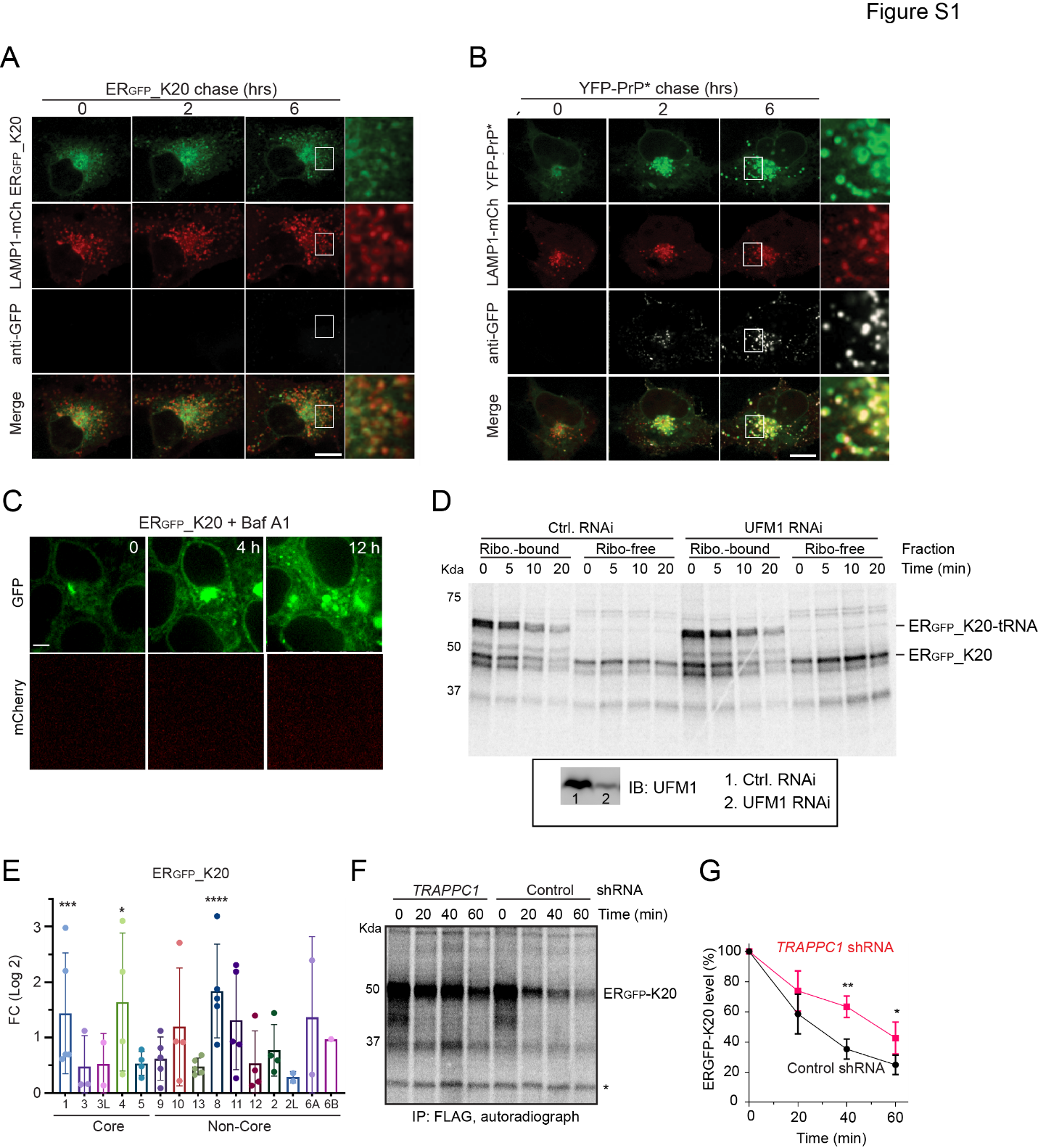

**Figure S1 The TRAPP complex promotes the transport of ERGFP_K20 to lysosomes via the Golgi. (Related to Figure 1)**

**(A, B)** As in Figure 1c except that the experiment was conducted in COS-7 cells transfected with ERGFP_K20 and LAMP1-mCherry (mCh) **(A)** or YFP-PrP* and LAMP1-mCherry **(B)**. Cells were imaged after treatment with Baf A1 (200 nM) and Alexa647-labeled anti-GFP antibodies. The representative images from time lapse videos show that ERGFP_K20 and YFP-PrP* are transported to lysosomes via distinct mechanisms. Scale bar, 10 μm.

**(C)** Baf A1 treatment did not increase the readthrough product of ERGFP_K20. HEK293T cells transfected with ERGFP_K20 were treated with Baf A1 and imaged. This panel serves as a control for Figure 1C. Scale bar, 5 μm.

**(D)** UFM1 is not required for substrate release from ribosome. Cells transfected with ERGFP_K20 were pulsed labeled with S^35^-Met/Cys and then incubated with unlabeled Met/Cys in excess for the indicated time point. Cell extracts were fractionated into a ribosome and a ribosome-free fraction for immunoprecipitation of ERGFP_K20. A fraction of the lysate was analyzed by immunoblotting to verify the knockdown efficiency.

**(E)** sgDNAs targeting the TRAPP complex subunits were enriched in the ERGFP_K20 (A) high cells. *, *p*<0.05; ***, *p*<0.005; ****, *p*<0.0001.

**(F, G)** TRAPPC1 depletion stabilizes ERGFP_K20. Pulse chase analysis of ERGFP_K20 turnover in control and TRAPPC1-depleted cells. The asterisk indicates a non-specific band that serves as a loading control. The graph in **(G)** shows the quantification of the experiment. Error bars indicate means ± SD; *, *p*<0.05; **, *p*<0.01 by unpaired Student *t*-test, n=3.

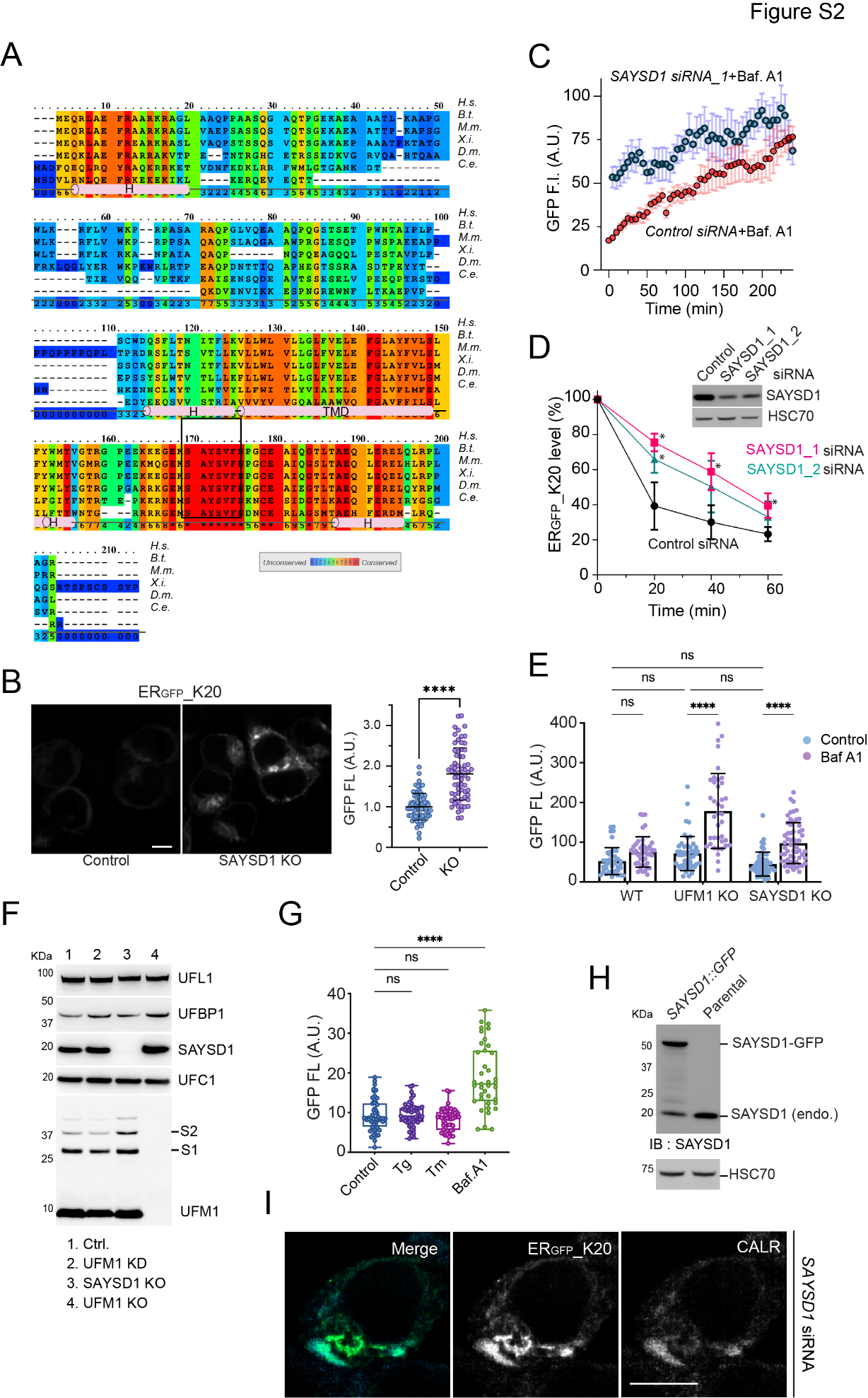

**Figure S2 SAYSD1 promotes the degradation of ERGFP_K20. (Related to Figure 2)**

**(A)** The sequence alignment and predicted secondary structure of SAYSD1 homologues. H, helix; TMD, transmembrane domain. The box indicates the highly conserved SAYS_V_FN motif. *H.s., Human; B.t., Cow; M.m., Mouse; X.i., Frog; D.m., Fly; C.e., Worm.*

**(B)** CRISPR-mediated knockout (KO) of SAYSD1 causes accumulation of ERGFP_K20. Scale bar, 5 µm. The graph shows the quantification of the fluorescence intensity of individual cells for GFP. Error bars indicate means ± SD ****; *p*<0.0001 by unpaired Student’s *t*-test.  n=3 independent experiments.

**(C)** Live-cell imaging analyses of the kinetics of ERGFP_K20 accumulation in cells treated with the indicated siRNA and Baf A1. Shown is the averaged GFP intensity from 8 randomly selected fields each with at least 4 cells. A.U. arbitrary unit. Error bars, mean ± SD, n=8.

**(D)** Pulse chase analysis of ERGFP_K20 turnover in ERGFP_K20 cells transfected with either control or *SAYSD1* siRNAs. The immunoblotting panels show the knockdown efficiency from a representative experiment. Error bars indicate means ± SD, * *p*<0.05 by unpaired Student’s *t*-test. n=3 independent experiments.

**(E)** SAYSD1 or UFM1 depletion does not cause accumulation of ER_GFP__K0. WT 293T cells or the indicated CRISPR KO cells were transfected with ER_GFP__K0 for 24 h followed by treatment with either DMSO control or Baf A1 (200nM) for 6 h. Cells were imaged and GFP fluorescence (FL) was quantified by Image J. Error bars, mean ± SD, **** p<0.0001 by unpaired Student’s t-test; ns, non-significant.

**(F)** SAYSD1 knockout (KO) does not reduce the expression of UFM1 or UFMylating enzymes. HEK293T cell lysates from the indicated genetic backgrounds were analyzed by immunoblotting. KD, knockdown.

**(G)** ER stress does not stabilize ER_GFP__K20. ER_GFP__K20 stable cells were treated with the indicated drugs for 6 h before imaging. Shown is the quantification of GFP intensity in at least 20 cells. Tg, Tharpsigargin (100nM); Tm, Tunicamycin (5 µg/mL). Error bars, mean ± SD, **** p<0.0001; ns, non-significant by unpaired Student’s t-test.

**(H)** Immunoblotting analysis confirms the correct insertion of a GFP-encoding sequence in the endogenous SAYSD1 locus.

**(I)** ERGFP_K20 accumulated in *SAYSD1* knockdown cells is co-localized with endogenous Calreticulin. Cells transfected with SAYSD1 siRNA were fixed and stained with Calreticulin antibody (CALR). Scale bar, 10 μm.

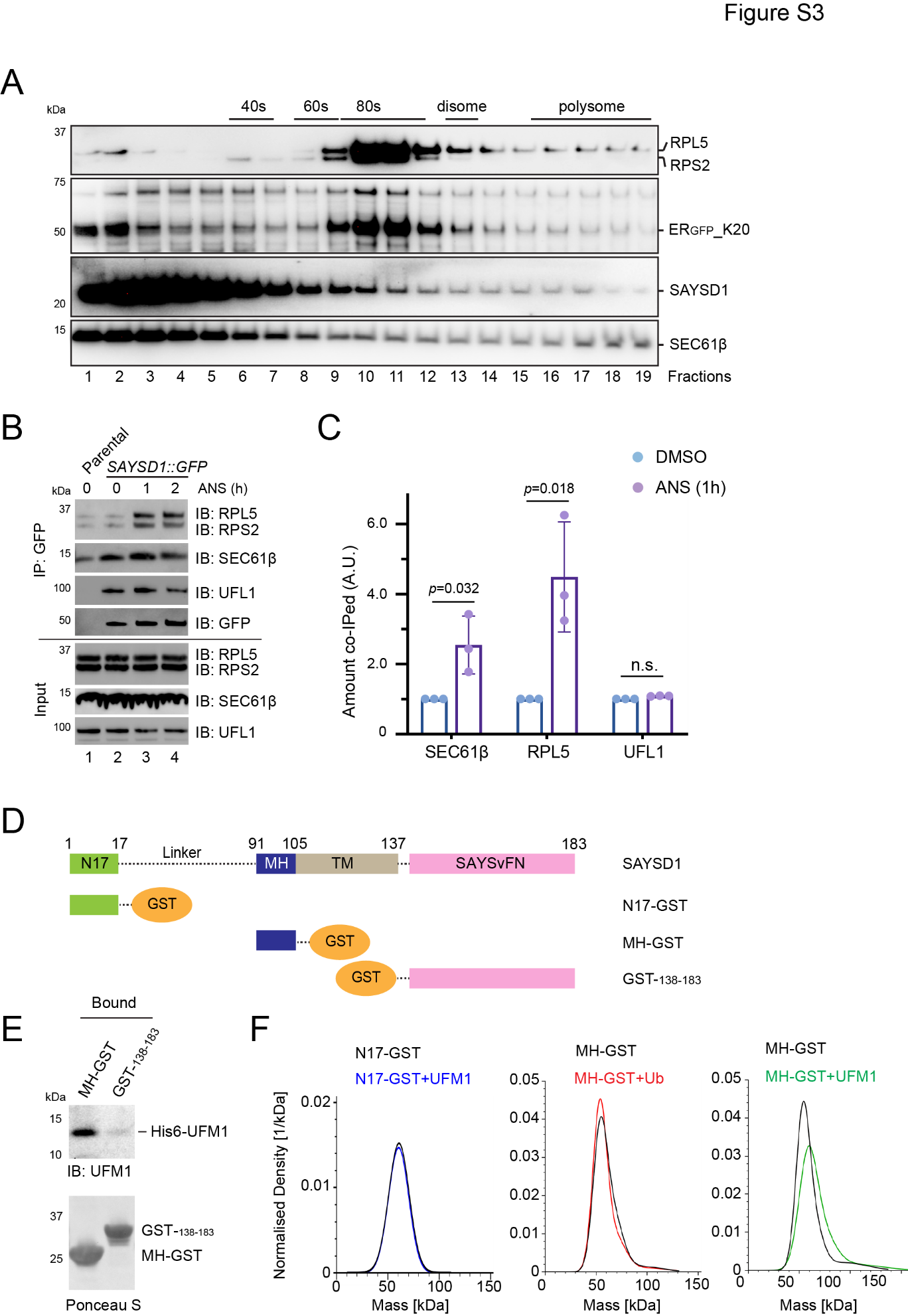

**Figure S3 Translation stalling stabilizes the interaction of SAYSD1 with Sec61 and ribosome.** **(Related to Figure 3)**

**(A)** SAYSD1 co-fractionates with Sec61β. Cell lysate prepared from 293T cells transfected with ER_GFP__K20 was subject to sucrose gradient centrifugation. Fractions were analyzed by immunoblotting.

**(B, C)** Proteins immunoprecipitated by GFP antibodies from *SAYSD1::GFP* cells treated with anisomycin (ANS, 200 nM) as indicated were analyzed by immunoblotting **(B)**. The graph in **(C)** shows the quantification of the normalized proteins co-precipitated with SAYSD1-GFP from cells treated with ANS for 1 h. Error bars indicate means ± SD; *p* values are from unpaired Student’s *t*-tests; n.s. not significant; n=3 independent experiments.

**(D)** A schematic diagram of the SAYSD1 domain structure and the recombinant SAYSD1 fragments tested in the study.

**(E)** GST pulldown shows that the middle helical (MH) segment in SAYSD1 but not the C-terminal domain binds UFM1.

**(F)** MH-GST binds specifically to UFM1. The indicated GST-tagged proteins (60nM) were mixed with either UFM1 or Ubiquitin (Ub) (120nM). The mixed samples were analyzed by mass photometry.

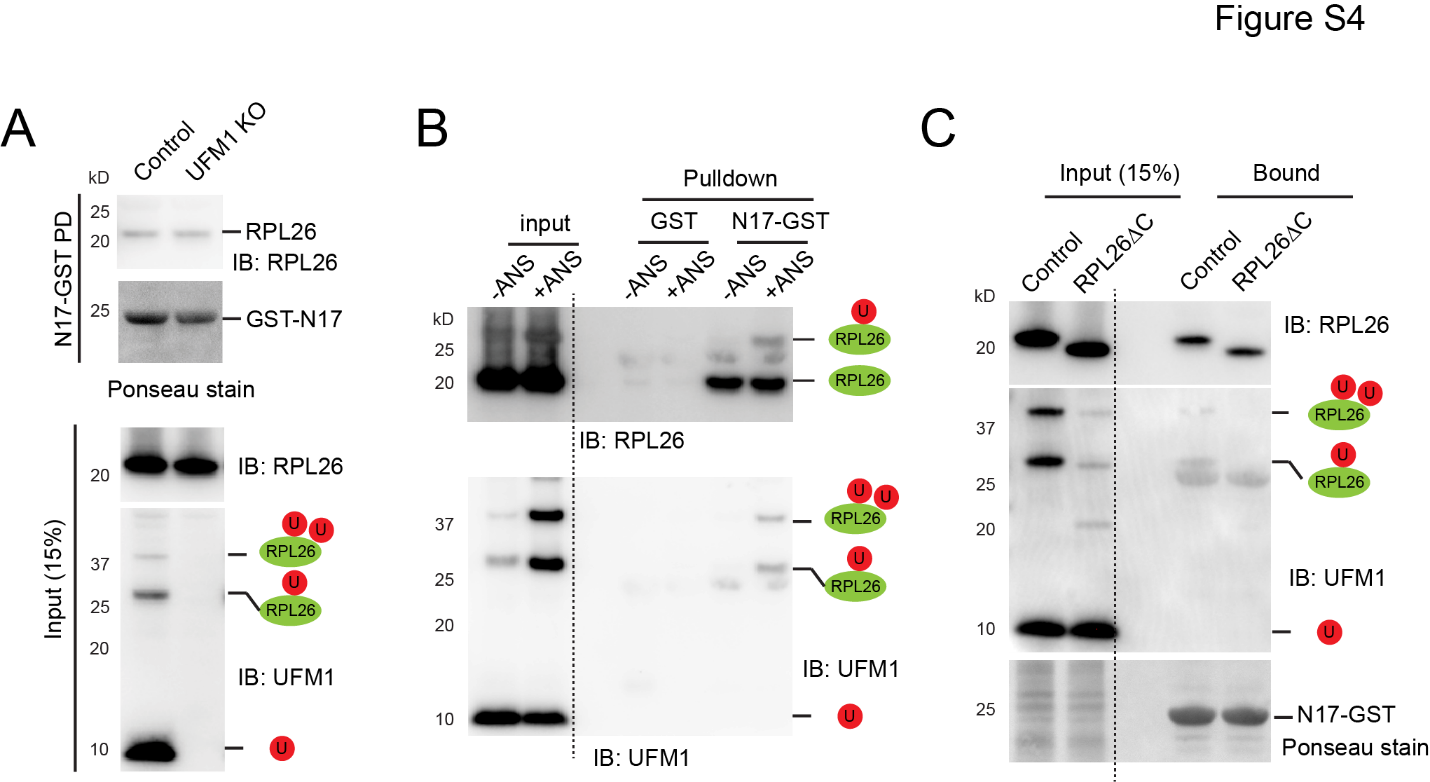

**Figure S4 The SAYSD1_N17 domain binds ribosome to promote TAQC. (Related to Figure 4)**

**(A)** UFM1 knockout (KO) does not affect the N17 and ribosome interaction. GST-N17 immobilized on glutathione beads was incubated with extracts from either wildtype control or UFM1 KO cells. Precipitated proteins were analyzed by immunoblotting. U stands for UFM1.

**(B)** GST or GST-N17 was immobilized on glutathione beads and then incubated with either control extract (-ANS) or an extract from ANS-treated HEK293T cells (+ANS). Proteins precipitated were analyzed by immunoblotting.

**(C)** GST-N17 immobilized on glutathione beads was incubated with extract from either control wildtype cells or cells that lack the C-terminal UFMylation site on endogenous RPL26.

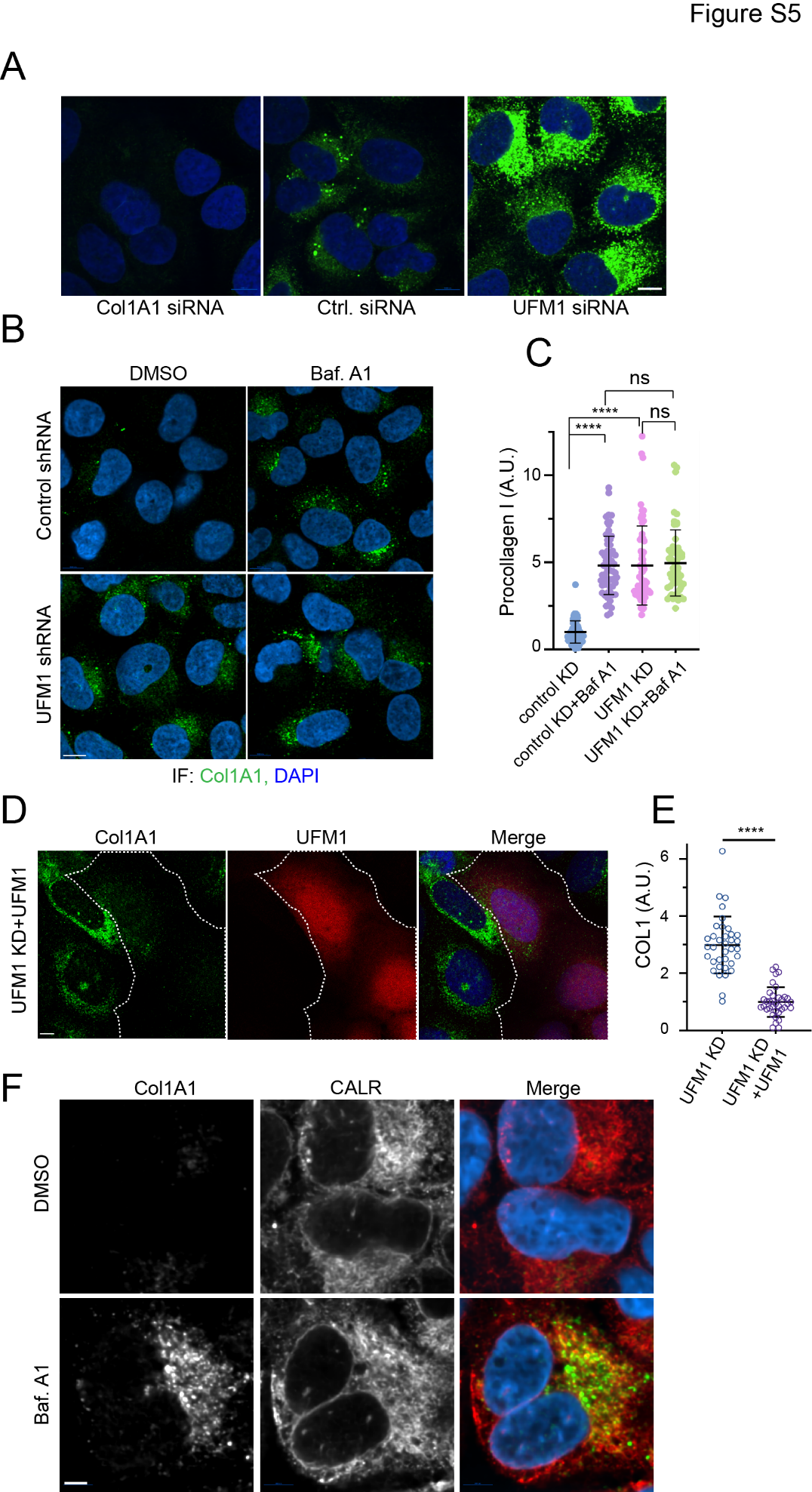

**Figure S5 Depletion of UFM1 or SAYSD1 causes accumulation of Col1A1 in mammalian cells. (Related to Figure 5)**

**(A-C)** UFM1 knockdown inhibits lysosome-mediated degradation of endogenous Col1A1. **(A)** U2OS cells treated with the indicated siRNA were stained by Col1A1 antibodies (Green) and DAPI (Blue). Scale bar, 10 µm. **(B)** Control and UFM1 knockdown U2OS cells were treated with DMSO or with Baf A1 (200 nM, 5 h), and stained by Col1A1 antibodies (Green) and DAPI (Blue). Scale bar, 10 µm. **(C)** Quantification of the Col1A1 signal in experiments represented by **(B)**. Error bars, means ± SD; ****, p-value<0.0001, by one-way ANOVA followed by Dunnett’s multiple comparisons test. n=3 independent experiments.

**(D, E)** UFM1 re-expression restores the low levels of Col1A1 in UFM1 knockdown cells. **(D)** UFM1 knockdown cells were transfected with a UFM1-expressing plasmid. Cells were stained by Col1A1 (green) and UFM1 (red) antibodies. The graph in **E** shows the quantification of Col1A1 fluorescence intensity in individual cells. A.U., arbitrary units. Error bars, means ± SD; ****, *p*<0.0001 by unpaired Student’s *t*-test. n=3. Scale bar, 5 µm.

**(F)** Baf A1 treatment stabilizes Col1A1 in a post-ER compartment. U2OS cells treated with DMSO as a control or with Baf A1 (250 nM, 5 h) were stained by Col1A1 and Calreticulin (CALR) antibodies. Scale bars, 5 μm.

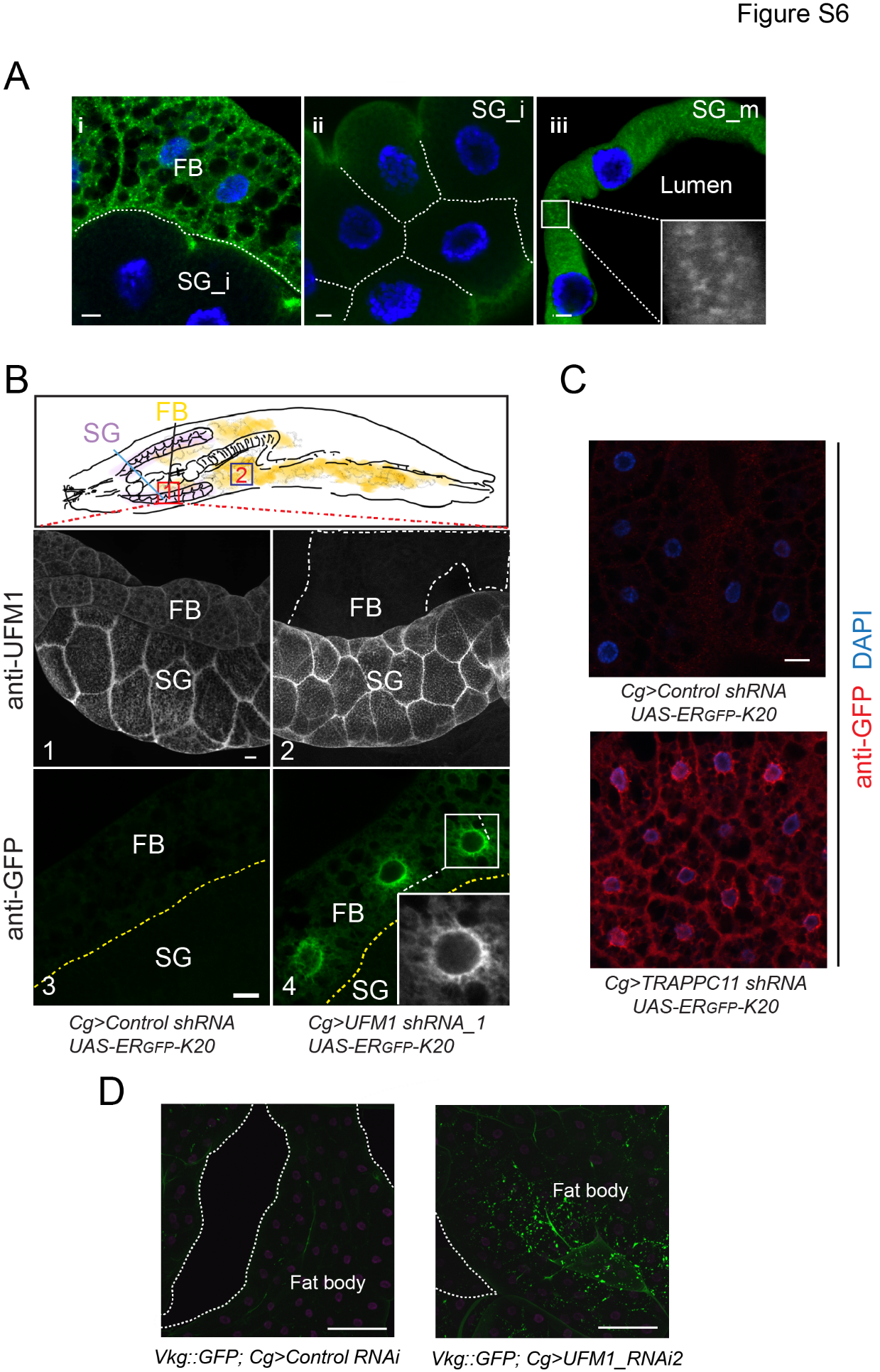

**Figure S6 UFM1- and SAYSD1-mediated TAQC is required for Col4 quality control in *Drosophila.* (Related to Figure 6)**

**(A)** UFM1 is highly expressed in secretory tissues in flies. The expression of UFM1 in *Drosophila* larval salivary gland (SG) or SG-associated fat body (FB) were analyzed by immunostaining with a UFM1 antibody (green) and DAPI (blue). SG_i, immature SG; SG_m, mature SG. Scale bar, 10 µm.

**(B, C)** Fat body (FB) specific knockdown of UFM1 **(B)** or TRAPPC11 **(C)** by the Cg-gal4 driver causes accumulation of ERGFP_K20. The diagram in **B** (top panel) indicates the location of the SG-associated FB (box 1) analyzed in **B** or gut-associated FB (box 2) in **C**. For *UFM1* knockdown, anti-UFM1 antibody staining in **B** (panels 1, 2) verifies FB-specific UFM1 depletion. GFP antibody staining detects perinuclear accumulation of ERGFP_K20 in UFM1 knockdown FB (panel 4 vs. 3). For TRAPPC11 knockdown **(c)**, the tissues were stained with GFP antibodies (red) and DAPI (blue). Scale bars, 20 µm.

**(D)** FB specific depletion of UFM1 using a second shRNA-expressing line also causes accumulation of Viking-GFP (Vkg-GFP). The tissues were stained with DAPI in magenta to reveal the nuclei. Scale bars, 100 μm.

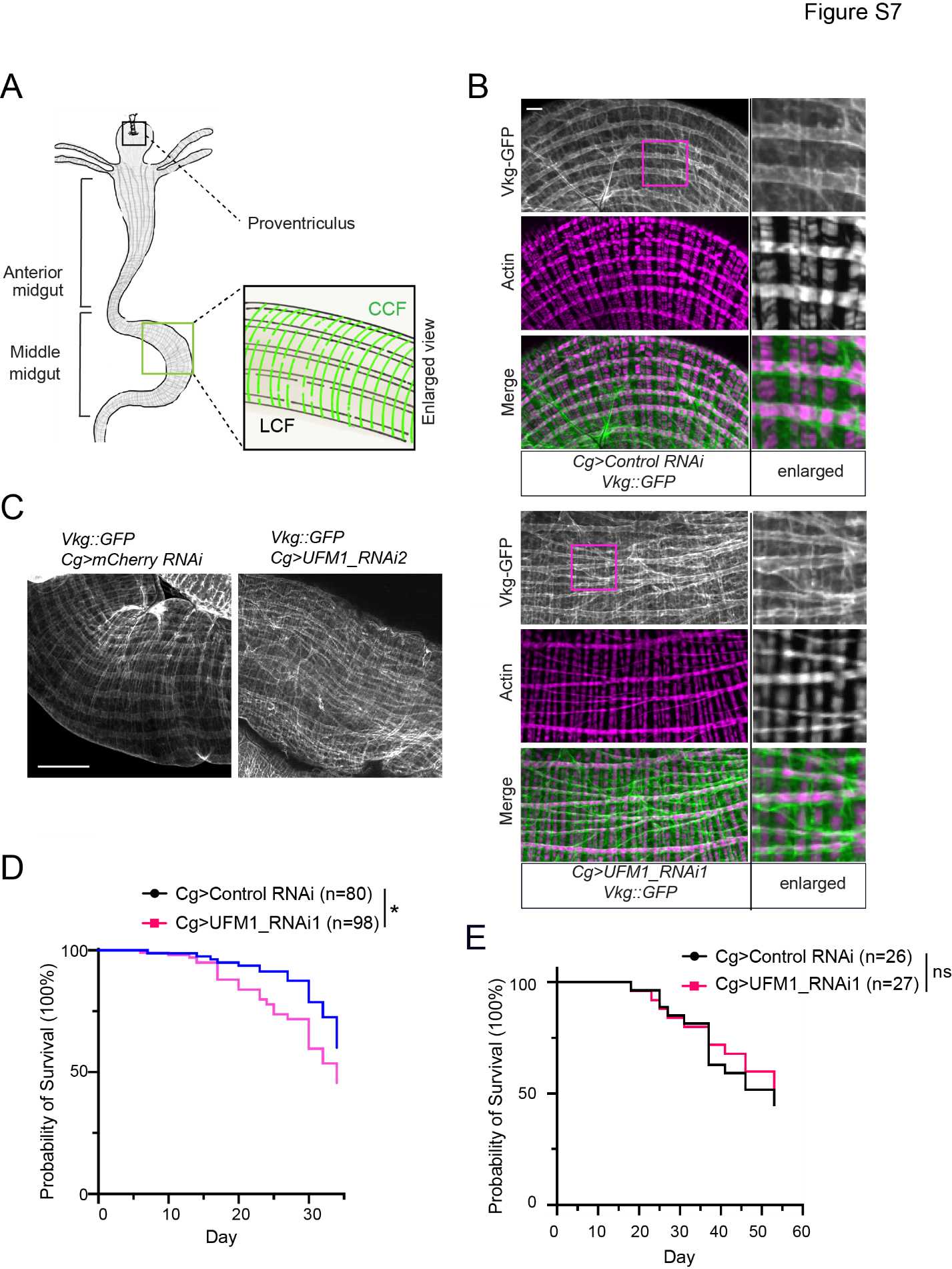

**Figure S7 UFM1 deficiency causes abnormal Col4 deposition. (Related to Figure 7)**

**(A)** A schematic diagram of the larval gut system. The regions chosen for analyses in Fig. 7 were indicated by the boxes.

**(B)** FB specific depletion of UFM1 disrupts the Vkg-GFP-containing collagen fibril pattern on middle midgut in third instar larvae. Scale bars, 100 μm.

**(C)** Abnormal Viking-containing basement membranes on middle midgut of adult flies with FB specific depletion of UFM1. The guts from adult females of the indicated genotypes were stained by phalloidin to label cortex actin (magenta). Small panels show enlarged views of the box-indicated areas. Scale bar, 20 µm.

**(D)** FB specific knockdown of UFM1 reduces the life span of flies at a high temperature. Flies of the indicated genotypes were raised at 29 °C and scored for viability. **(E)** Same as **D** except that the flies were raised at 25 °C. n indicates fly numbers from two independent experiments. *, *p*=0.011, by Gehan-Breslow-Wilcoxon test.
